# Axonal organelle buildup from loss of AP-4 complex function causes exacerbation of amyloid plaque pathology and gliosis in Alzheimer’s disease mouse model

**DOI:** 10.1101/2024.03.31.587499

**Authors:** Alex Orlowski, Joseph Karippaparambil, Jean-Michel Paumier, Shraddha Ghanta, Eduardo Pallares, Rumamol Chandran, Daisy Edmison, Jamuna Tandukar, Ruixuan Gao, Swetha Gowrishankar

## Abstract

Lysosomes and related precursor organelles robustly build up in swollen axons that surround amyloid plaques and disrupted axonal lysosome transport has been implicated in worsening Alzheimer’s pathology. Our prior studies have revealed that loss of Adaptor protein-4 (AP-4) complex function, linked primarily to Spastic Paraplegia (HSP), leads to a similar build of lysosomes in structures we term “AP-4 dystrophies”. Surprisingly, these AP-4 dystrophies were also characterized by enrichment of components of APP processing machinery, β-site cleaving enzyme 1 (BACE1) and Presenilin 2. Our studies examining whether the abnormal axonal lysosome build up resulting from AP-4 loss could lead to amyloidogenesis revealed that the loss of AP-4 complex function in an Alzheimer’s disease model resulted in a strong increase in size and abundance of amyloid plaques in the hippocampus and corpus callosum as well as increased microglial association with the plaques. Interestingly, we found a further increase in enrichment of the secretase, BACE1, in the axonal swellings of the plaques of Alzheimer model mice lacking AP-4 complex compared to those having normal AP-4 complex function, suggestive of increased amyloidogenic processing under this condition. Additionally, the exacerbation of plaque pathology was region-specific as it did not increase in the cortex. The burden of the AP-4 linked axonal dystrophies/AP-4 dystrophies was higher in the corpus callosum and hippocampus compared to the cortex, establishing the critical role of AP-4 -dependent axonal lysosome transport and maturation in regulating amyloidogenic amyloid precursor protein processing.

**Significance Statement:** A major pathological feature of Alzheimer’s disease is the accumulation of axonal lysosomes near sites of amyloid plaques. Lysosome accumulation is thought to contribute to amyloid production. In fact, a genetic perturbation that arrests lysosomes in axons exacerbates amyloid plaque pathology. The mechanisms that control axonal lysosome abundance as well the molecular composition of axonal endolysosomes that produce Abeta, however, are not fully understood. Axonal lysosome build-up is emerging as a common pathology in other neurodegenerative disorders such as Hereditary Spastic Paraplegia (HSP), but its relevance to amyloid production is unknown. We find that a model of HSP caused by loss of AP-4 adaptor complex lead to axonal lysosome buildup that differs in some of its content, but still contributes to amyloidogenesis. This demonstrates that different perturbations leading to changes in heterogeneous pool of axonal lysosomes can converge on a common pathology.

## Introduction

Alzheimer’s disease (AD) is a devastating neurodegenerative disease that is also the leading cause of dementia among the elderly. A defining pathological feature of AD is the amyloid plaque or the neuritic plaque, which is composed of a core of extracellular aggregated amyloid β (Aβ), dystrophic/swollen axons and microglia (Terry et al., 1964; Itagaki et al., 1989; Fiala, 2007; Condello et al., 2011). The dystrophic axons themselves contain massive accumulations of lysosome-related organelles including late endosomal and lysosomal intermediates, autophagosomes as well as hybrid organelles arising from the fusion of autophagic and lysosomal pathway intermediates (Terry et al., 1964; Su et al., 1993; Cataldo et al., 1994; Nixon, 2005; Condello et al., 2011; Gowrishankar et al., 2015). Investigations into the nature and disease relevance of these organelle accumulations to AD pathology seems to suggest that the accumulating lysosomes could contribute to the amyloid pathology. Studies carried out on human brain tissue as well as transgenic mouse models of AD have revealed that the swollen axons around Aβ aggregates are enriched in the amyloid precursor protein (APP) and BACE1, the β-secretase involved in the amyloidogenic processing of APP (Cras et al., 1991; Cummings et al., 1992; Kandalepas et al., 2013; Yoon et al., 2013; Gowrishankar et al., 2015). Observations from human AD brain support the presence of PSEN1, a component of the γ−secretase complex, in lysosome-like organelles at swollen axons of neuritic plaques (Yu et al., 2005). Furthermore, PSEN2, the PSEN that constitutively localizes to lysosomes (Sannerud et al., 2016) was also observed to be highly enriched in axonal swellings at amyloid plaques of transgenic AD mice (Gowrishankar et al., 2017). Importantly, depleting JIP3, a lysosomal adaptor that is critical for optimal retrograde axonal lysosome transport (Drerup and Nechiporuk, 2013; Edwards et al., 2013; Gowrishankar et al., 2017), from 5xFAD mice resulted in worsening of amyloid plaque pathology (Gowrishankar et al., 2017). In further support of a model where lysosomes accumulating in swollen axons could contribute to Aβ production, loss of JIP3/ MAPK8IP3 protein from primary mouse cortical neurons as well as human iPSC derived neurons (i^3^Neurons) was sufficient to cause both an axonal lysosome build up as well as increased intraneuronal Aβ42 (Gowrishankar et al., 2017; Gowrishankar et al., 2021). Recent studies have also reported the enrichment of the pre-autophagosomal protein ATG9A in dystrophic axons around amyloid plaques (Sharoar et al., 2019; Edmison et al., 2021). Interestingly, loss of the adaptor protein complex -4 (AP-4), the complex responsible for correct sorting of ATG9 out of the trans-golgi network (Mattera et al., 2017; Davies et al., 2018), also results in accumulation of lysosomes (LAMP1-positive compartments) in axonal swellings (Mattera et al., 2017; Edmison et al., 2021; Majumder et al., 2022). Consistent with the model of endo-lysosomal intermediates being sites of APP processing and Aβ production, the AP-4ε KO mice exhibit enrichment of the β-secretase, BACE1, and subunit of the γ-secretase complex, PSEN2, in their axonal dystrophies (Edmison et al., 2021). While the AP-4ε KO condition closely resembled JIP3 loss of function in terms of the accumulation of lysosomes and enrichment of BACE1 and PSEN2, our prior studies demonstrated that there was a surprising enrichment of JIP3 in axonal swellings in human i^3^Neurons with reduced AP-4 complex function (Majumder et al., 2022). This unexpected finding suggests that there are likely differences between the nature of lysosomes accumulating under these two conditions including their immediate environment: for instance, JIP3 levels in the axoplasm around them. To determine if the axonal lysosomes building up in the AP-4ε KO mouse brain could contribute to amyloidogenic processing of APP, despite potential differences from those organelles observed previously from JIP3 loss of function, we generated 5xFAD animals lacking the AP-4ε subunit (and thus functional AP-4 complex; (De Pace et al., 2018)), and examined the development of plaque pathology under this condition. We found that the loss of AP-4 complex strongly exacerbated amyloid plaque pathology, including abundance and size of amyloid plaques in the corpus callosum and hippocampus. However, this worsening of plaque pathology was not observed in the cortex. Interestingly, this exacerbation correlated with where AP-4 dependent axonal dystrophies (which we term “AP-4 dystrophies”) are strongest in AP-4ε KO animals, with the corpus callosum and hippocampus having a significantly higher burden than the cortex. In addition to the increased size of the neuritic plaques, we also observed an increase in microglial recruitment to these plaques in the absence of AP-4 complex. 5xFAD animals lacking AP-4 complex continued to exhibit AP-4 dystrophies in addition to the neuritic plaques and were readily distinguishable from the neuritic plaques by their morphology, relatively smaller size and the lack of extracellular Aβ aggregate in their immediate vicinity. Quantitative measurements of enrichment of the proteins revealed that BACE1 was more enriched in the dystrophic axons of neuritic plaques in the 5xFAD; AP-4ε KO animals compared to those in 5xFAD; AP-4 WT animals. Our studies here suggest that increased axonal lysosome abundance by distinct mechanisms can contribute to amyloidogenic APP processing so long as there is a consequent increase in APP processing machinery in these organelles.

## Results

### Loss of AP-4 complex function increases amyloid plaque pathology

The presence of lysosome-filled axonal swellings is a major pathological feature of Alzheimer’s disease amyloid plaques. Studies that perturbed axonal lysosome transport suggest that these organelles can contribute to amyloidogenic processing of APP and consequent worsening of plaque pathology. In recent years, several other conditions have also been shown to cause lysosomes and lysosome-related organelles to build up in axonal swellings (Edmison et al., 2021; Roney et al., 2022). Our prior studies on the nature and composition of such organelles that build up from loss of AP-4 complex function indicated that critical components of the APP processing machinery also build up here (Edmison et al., 2021). To determine if these AP-4 linked axonal dystrophies, referred to henceforth as AP-4 dystrophies, and resultant organelle build up contribute to Alzheimer’s disease pathology, we carried out a comparative analysis of amyloid plaque pathology and gliosis in the 5xFAD mice (Oakley et al., 2006) that carried normal amounts of the AP-4ε subunit (5xFAD; AP-4 WT) with 5xFAD mice lacking the AP-4ε subunit (5xFAD; AP-4 KO). Since the AP-4 complex is an obligate heterotetramer (Dell’Angelica et al., 1999; Hirst et al., 1999), the homozygous loss of the ε subunit of AP-4, renders the entire complex unstable and non-functional (De Pace et al., 2018; Ivankovic et al., 2020). A systematic analysis of the size of the neuritic plaques present in the hippocampus and corpus callosum as measured by area of amyloid deposits (Aβ staining) and lysosome-filled dystrophic axons (LAMP1 staining) that surround the amyloid aggregates revealed that there was a substantial increase in the mean individual size of the plaques upon loss of AP-4 complex (Figure 1A-G). Indeed, there was a ∼two-fold increase in mean size of both the amyloid aggregate (Figure 1D) and axonal dystrophies (Figure 1F). In addition to the increase in size of the individual plaques, there was a trend towards increase in the total number of plaques in these brain regions, upon loss of the AP-4 complex (Figure 1E, G; Figure S1). Examination of the correlation of size of amyloid deposits and lysosome filled dystrophic axons revealed that there was a positive relationship between the two for both genotypes (Figure 1H, I). However, the exacerbation upon AP-4 loss is apparent from the fact that a fourth of the neuritic plaques in the 5xFAD; AP-4 KO animals were much bigger than any observed in the 5xFAD; AP-4 WT animals (Figure 1H, I). A trend towards increased plaque pathology was also observed in male 5xFAD; AP-4 KO mice when compared to 5xFAD; AP-4 WT mice (Figure S2). This exacerbation of amyloid plaque pathology was region-specific as the plaque size and abundance measured in the cerebral cortex did not show an increase upon loss of AP-4 complex, but in fact a slight trend towards decrease (Figure S3). Thus, loss of AP-4 complex resulted in an increase in amyloid plaque pathology specifically in the corpus callosum and hippocampal regions.

**Figure 1:**
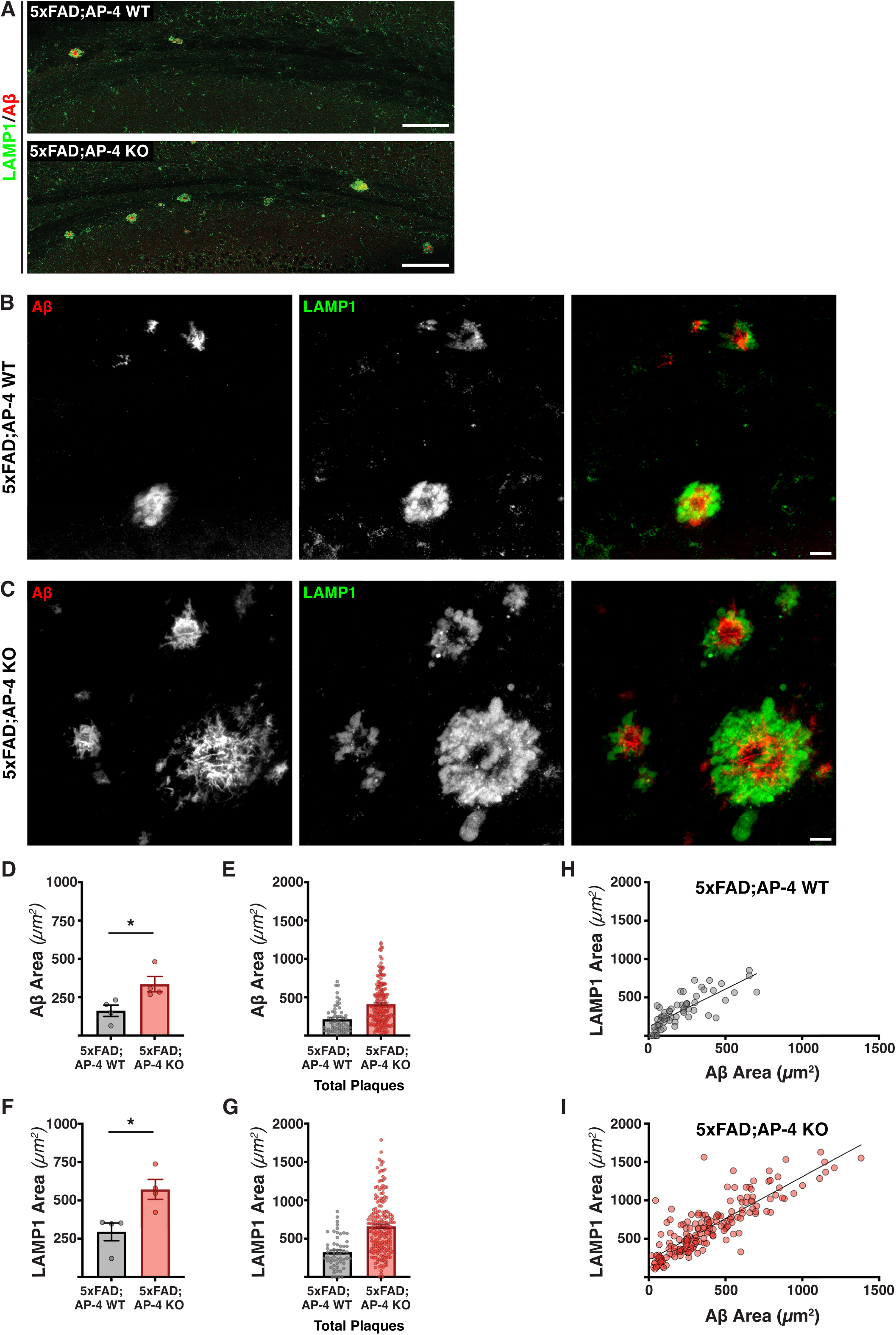
Loss of AP-4 complex enhances amyloid plaque pathology in 3-month-old 5xFAD mice. (A) Stitched confocal images of corpus callosum of 3-month-old 5xFAD mice with AP-4 (WT) or lacking AP-4 (KO) stained for LAMP1 (green; lysosomes) and Aβ (red; amyloid aggregates) depicting neuritic plaques. Bar, 100μm. (B and C) High-magnification confocal images of neuritic plaques labeled with LAMP1 (green; lysosomes) and Aβ (red; amyloid aggregates) in 5xFAD; AP-4 WT (B) and 5xFAD; AP-4 KO (C). Bar, 10μm. (D) Quantification of mean size of amyloid aggregates of neuritic plaques in 3-month-old female 5xFAD; AP-4 WT and KO mice (Grey-AP-4 WT; Red-AP-4 KO). Mean ± SEM, N = 4 pairs of 5xFAD; AP-4 WT and 5xFAD; AP-4 KO animals; *, P < .05, unpaired *t* test. (E) Plot of areas of each individual amyloid aggregate present in the hippocampus and corpus callosum of the four animals from each genotype (Grey-5xFAD; AP-4 WT; Red-5xFAD; AP-4 KO). (F) Quantification of mean size of lysosome-filled axonal swellings of neuritic plaques in 3-month-old female 5xFAD; AP-4 WT and KO mice (Grey-5xFAD; AP-4 WT; Red-5xFAD; AP-4 KO). Mean ± SEM, N = 4, *, P < .05, unpaired *t* test. (G) Plot of the areas of individual lysosome-filled axonal swellings present in the hippocampus and corpus callosum of the four animals from each genotype (Grey-5xFAD; AP-4 WT; Red-5xFAD; AP-4 KO). (H and I) Plot depicting the correlation between LAMP1 and Aβ areas of all individual neuritic plaques present in the hippocampus and corpus callosum of the four animals from each genotype-5xFAD; AP-4 WT (H) and 5xFAD; AP-4 KO animals (I).

### Loss of AP-4 complex results in increased microgliosis in Alzheimer’s disease

Having observed a substantial worsening in amyloid plaque pathology in 3-month-old 5xFAD; AP-4 KO animals, we next examined if there were any changes in microgliosis in these animals compared to 3-month-old 5xFAD; AP-4 WT mice. Microglial recruitment to amyloid plaques is thought to be an inflammatory response that occurs concurrently to the amyloid deposition (Song and Colonna, 2018; Leng and Edison, 2021). Given this, we examined the distribution as well as plaque-association of microglia in the corpus callosum and hippocampi of these mice by immunostaining for iba1 (a calcium binding protein in microglia) along with LAMP1 (labeling the dystrophic axons of neuritic plaques). While the stitched images obtained even at lower magnification (Figure 2A; Figure S4) revealed the marked increase in amyloid pathology as well as gliosis, we imaged all the individual plaques and associated glia within the hippocampus and corpus callosum at high magnification and resolution (Figure 2B, C). Our analysis of these plaques and associated microglia revealed that most neuritic plaques (68% of them in 5xFAD; AP-4 WT and 85% of plaques in 5xFAD; AP-4 KO animals) have one or more microglia associated with them (Figure 2D, E). A systematic analysis of size of each individual neuritic plaque as a function of the number of microglia associated with them (Figure 2F-I), revealed several interesting differences between 5xFAD animals with and without AP-4 complex. Firstly, several plaques in the 3-month-old 5xFAD; AP-4 KO had 5-10 microglia associated with them (Figure 2I), which is never observed in the 3-month-old 5xFAD animal (Figure 2H). In fact, 25% of all plaques have more than 3 microglia associated with them in the 5xFAD; AP-4 KO animals while only approximately 2.6% percentage of plaques in the 5xFAD; AP-4 WT animals do so (Figure 2F). Interestingly, the increased glial recruitment in the 5xFAD; AP-4 KO animals was not associated only with the very large plaques. A substantial fraction (∼30%) of even moderately sized plaques that were comparable in size to several similar plaques in control animals had three or more glia associated with them (Figure 2G).

**Figure 2:**
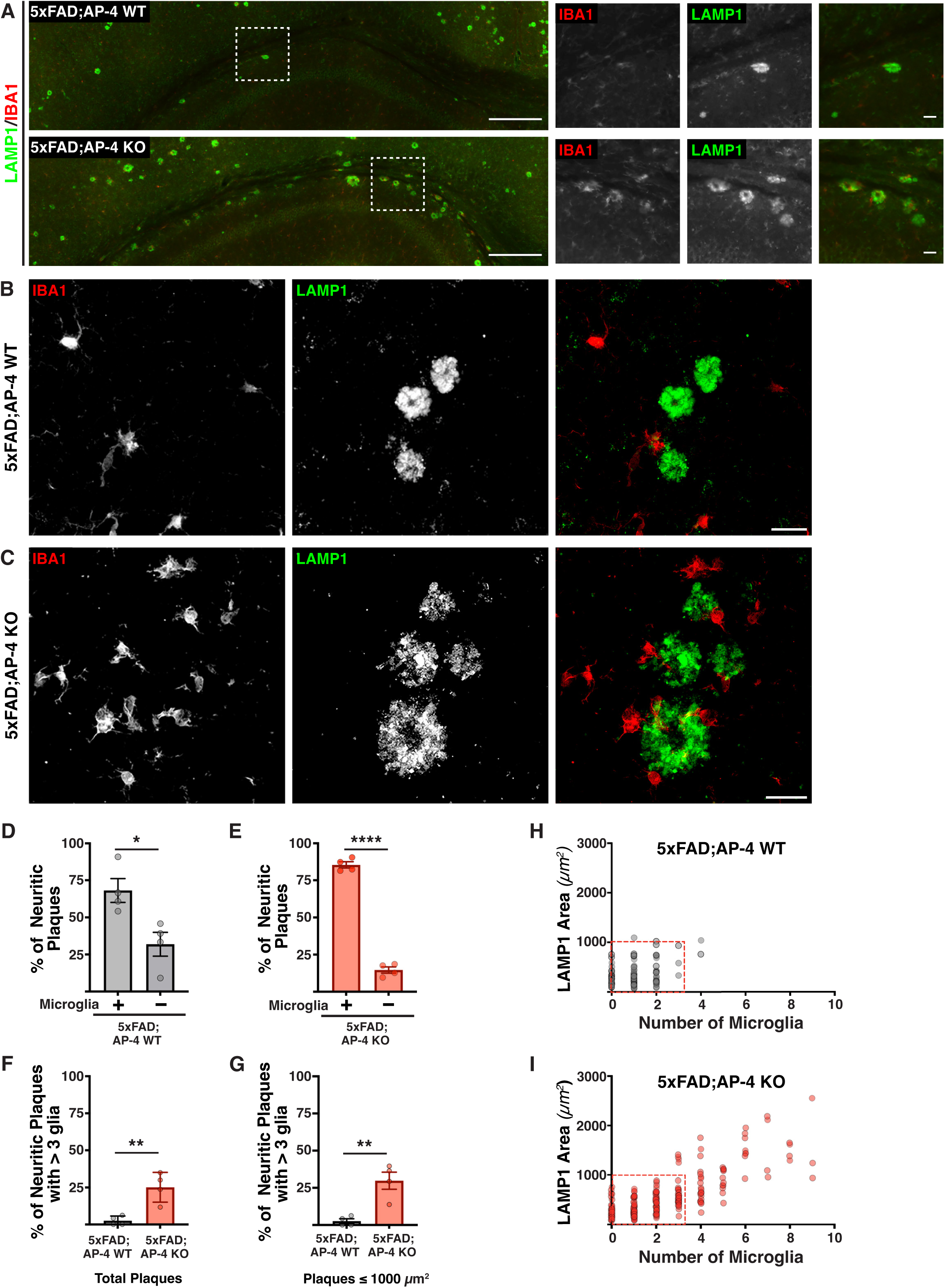
Loss of AP-4 complex leads to increased gliosis in 3-month-old 5xFAD mice. (A) Stitched images of corpus callosum and hippocampus of 3-month-old 5xFAD mice with AP-4 (WT) or lacking AP-4 (KO) depicting neuritic plaques stained with LAMP1 (green; lysosomes) and microglia stained with IBA1 (red). Bar, 100μm. The dashed white boxes in the stitched image are expanded in the inset images on the right of each stitched image. Bar, 10μm. (B and C) High-magnification images of neuritic plaques labeled with LAMP1 (green) and IBA1 (red) in 5xFAD; AP-4 WT (B) and 5xFAD; AP-4 KO (C). Bars, 10μm. (D) Quantification of percentage of neuritic plaques in 3-month-old female 5xFAD; AP-4 WT animals that have microglia associated with them. Mean ± SEM, N = 4, *, P < .05, unpaired *t* test. (E) Quantification of percentage of neuritic plaques in 3-month-old female 5xFAD; AP-4 KO animals that have microglia associated with them. Mean ± SEM, N = 4, ****, P < 0.0001, unpaired *t* test. (F) Percentage of total neuritic plaques in the hippocampus and corpus callosum of 3-month-old female 5xFAD; AP-4 WT (grey) and 5xFAD; AP-4 KO (red) animals with more than three microglia associated with them. Mean ± SEM, N = 4, **, P < 0.01, unpaired *t* test. (G) Percentage of moderately sized neuritic plaques (with areas under 1000μm^2^) in the hippocampus and corpus callosum of 3-month-old female 5xFAD; AP-4 WT (grey) and 5xFAD; AP-4 KO (red) animals with more than three microglia associated with them. Mean ± SEM, N = 4, **, P < 0.01, unpaired *t* test. (H and I) Plot depicting the correlation between neuritic plaque area and the number of associated microglia for all individual neuritic plaques present in the hippocampus and corpus callosum of the four animals from each genotype: 5xFAD; AP-4 WT (H) and 5xFAD; AP-4 KO mice (I).

### AP-4 dystrophies in Alzheimer’s disease mice differ from neuritic plaques in their size, glial association and amyloid deposition

Since our prior observation of the AP-4 dystrophies carrying organelles enriched in APP processing machinery had motivated us to examine their contribution to amyloid plaque pathology, we next examined if these dystrophies persisted in the 5xFAD background. Indeed, high resolution imaging of LAMP1 and Aβ staining (Figure 3A; Figure S5) revealed that in addition to amyloid/neuritic plaques, the 5xFAD; AP-4 KO animals also exhibited these AP-4 dystrophies (Figure 3A; yellow arrowhead; Figure S5) that were never observed in 5xFAD animals with normal AP-4 complex function/levels. Studies that examined the size of these individual dystrophies as well as those of the neuritic plaques in the same brain regions of these animals also revealed that they were considerably smaller than the neuritic plaques (Figure 3B). While differing in size, AP-4 dystrophies are highly similar to the dystrophies of neuritic plaques (Gowrishankar et al., 2015), in terms of endo-lysosomal organelles enriched in them. They are both enriched in LAMP1, positive for luminal lysosomal protein Progranulin, small GTPase RagC and relatively deficient in some soluble proteases such as Cathepsin B and Legumain (Figure S6). Interestingly, unlike the neuritic plaques that contained lysosome-filled axonal dystrophies surrounding the amyloid deposits, these AP-4 dystrophies did not appear to have extracellular amyloid aggregated detectable near them (Figure 3A, C), which we confirmed imaging at high resolution using expansion microscopy (Videos 1-3; Figure S5). A comparative analysis of all the individual neuritic plaques and the AP-4 dystrophies in terms of their sizes, of both associated amyloid deposit (Aβ staining) and axonal dystrophies (LAMP1 staining) in 5xFAD; AP-4 KO animals revealed that none of the AP-4 dystrophies (Figure 3C; Blue triangles) had any amyloid deposit in their immediate proximity, unlike the neuritic plaques (Figure 3C; Red circles). We then examined if these AP-4 dystrophies were associated with microglia. Consistent with our previous observations from AP-4ε KO animals, the majority (∼90%) of AP-4 dystrophies in 5xFAD; AP-4 KO animals did not have microglia associated with them (Figure 3D, E). Interestingly, in these 5xFAD, AP-4 KO animals, the size of AP-4 dystrophies that have microglia associated with them is significantly higher than those that do not (Figure 3F). Given the AP-4 dystrophies are yet to show amyloid deposition in their vicinity, they could represent an earlier stage in formation of the neuritic plaque. Consistent with this, the corpus callosum and hippocampus regions of AP-4ε KO animals exhibit a significantly higher (almost three-fold) burden of AP-4 dystrophies compared to the cortex (Figure 3G). Thus, the AP-4 dystrophy pathology arising from loss of AP-4 complex, is significantly higher in the same region where the amyloid plaque pathology is exacerbated in 5xFAD, AP-4 KO animals. Interestingly, loss of AP-4 complex by itself does not appear to cause increased gliosis, with AP-4ε KO animals showing no increase in gliosis in the corpus callosum and hippocampus regions, when compared to their wild type littermates (Figure S7).

**Figure 3:**
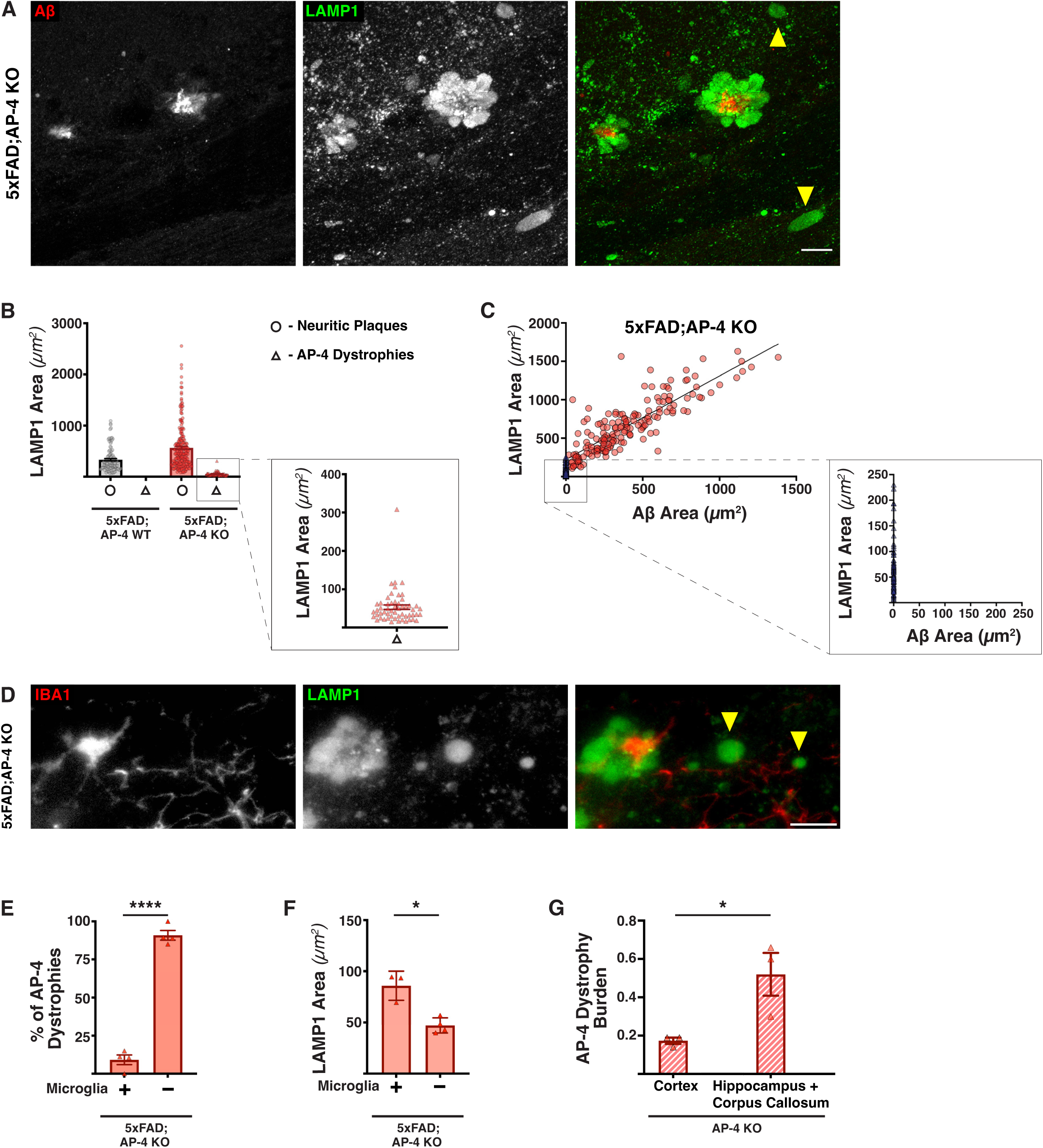
AP-4 axonal dystrophies observed in 5xFAD; AP-4 KO animals lack amyloid accumulation in their proximity. (A) High-magnification confocal images of LAMP1 (green; lysosomes) and Aβ (red; amyloid aggregate) in 5xFAD; AP-4 KO animals depicting both neuritic plaques and AP-4 dystrophies. Bar, 10μm. Yellow arrowheads point to AP-4 dystrophies. (B) Plot depicting the areas of all axonal dystrophies obtained from 5xFAD; AP-4 WT and 5xFAD; AP-4 KO animals; N = 4 animals per genotype. Neuritic plaques-circle and AP-4 dystrophies-triangle. (C) Plot depicting the correlation between LAMP1 and Aβ areas of all individual neuritic plaques and AP-4 dystrophies in 5xFAD; AP-4 KO animals; N = 4. Neuritic plaque-red circle; AP-4 dystrophy-blue triangle. (D) High-magnification images of LAMP1 (green; lysosomes) and IBA1 (red; microglia) in 5xFAD; AP-4 KO depicting neuritic plaques, AP-4 dystrophies and association of microglia. Bar, 10μm. Yellow arrowheads point to AP-4 dystrophies. (E) Quantification of percentage of AP-4 dystrophies that have microglia associated with them in the 3-month-old female 5xFAD; AP-4 KO animals. Mean ± SEM, N = 4, ****, P < 0.0001, unpaired *t* test. (F) Quantification of mean size of AP-4 dystrophies with and without microglia in 3-month-old female 5xFAD; AP-4 KO mice. Mean ± SEM, N = 4, *, P < .05, unpaired *t* test. (G) Quantification of AP-4 dystrophy burden (fraction of area occupied by AP-4 dystrophies out of total area) in corpus callosum, hippocampus versus cortex of 6-8-month-old AP-4ε KO animals. Mean ± SEM, N = 3, *, P < .05, unpaired *t* test.

### Loss of AP-4 complex leads to increased enrichment of BACE1 in the axonal swellings of the amyloid plaques

Our findings here that loss of AP-4 complex exacerbates amyloid plaque pathology including the size of amyloid deposits and abundance of these plaques along with our prior observations that AP-4 dystrophies themselves are also enriched in APP processing machinery led us to examine the relative enrichment of BACE1 in these axonal dystrophies. We examined the relative enrichment of both LAMP1 and BACE1 in the axonal dystrophies associated with neuritic plaques of both 5xFAD; AP-4 WT and 5xFAD; AP-4 KO as well as in AP-4 dystrophies in the latter (Figure 4A-C). We found that despite the larger size of neuritic plaques in the 5xFAD; AP-4 KO animals compared to neuritic plaques in the 5xFAD; AP-4 WT, they exhibited the same level of enrichment of the lysosomal membrane protein LAMP1 (Figure 4D, E; Figure S8A-C). In fact, this was comparable to the enrichment of LAMP1 in AP-4 dystrophies as well, despite these being far smaller on size (Figure 4D, E). However, we find that enrichment of BACE1 was higher in the axonal dystrophies of neuritic plaques in 5xFAD; AP-4 KO animals compared to those in 5xFAD; AP-4 WT, suggestive of potentially higher secretase activity in these organelles (Figure 4F, G; Figure S8A-B, D). Interestingly, while BACE1 is present in AP-4 dystrophies, its enrichment here is lower than in neuritic plaques present in either genotype (Figure 4F, G). While the AP-4 dystrophies appeared to resemble the dystrophies of neuritic plaques in terms of LAMP1 enrichment (Figure 4D, E) and have comparatively lesser BACE1enrichment (Figure 4F, G), we found that they have a significantly higher enrichment of the lysosomal GTPase, Arl8 (Figure S9). While the enrichment levels of Arl8 in neuritic plaques of both 5xFAD; AP-4 WT and 5xFAD; AP-4 KO were comparable, the enrichment of Arl8 in the AP-4 dystrophies is significantly higher than in neuritic plaques (Figure S9A-D).

**Figure 4:**
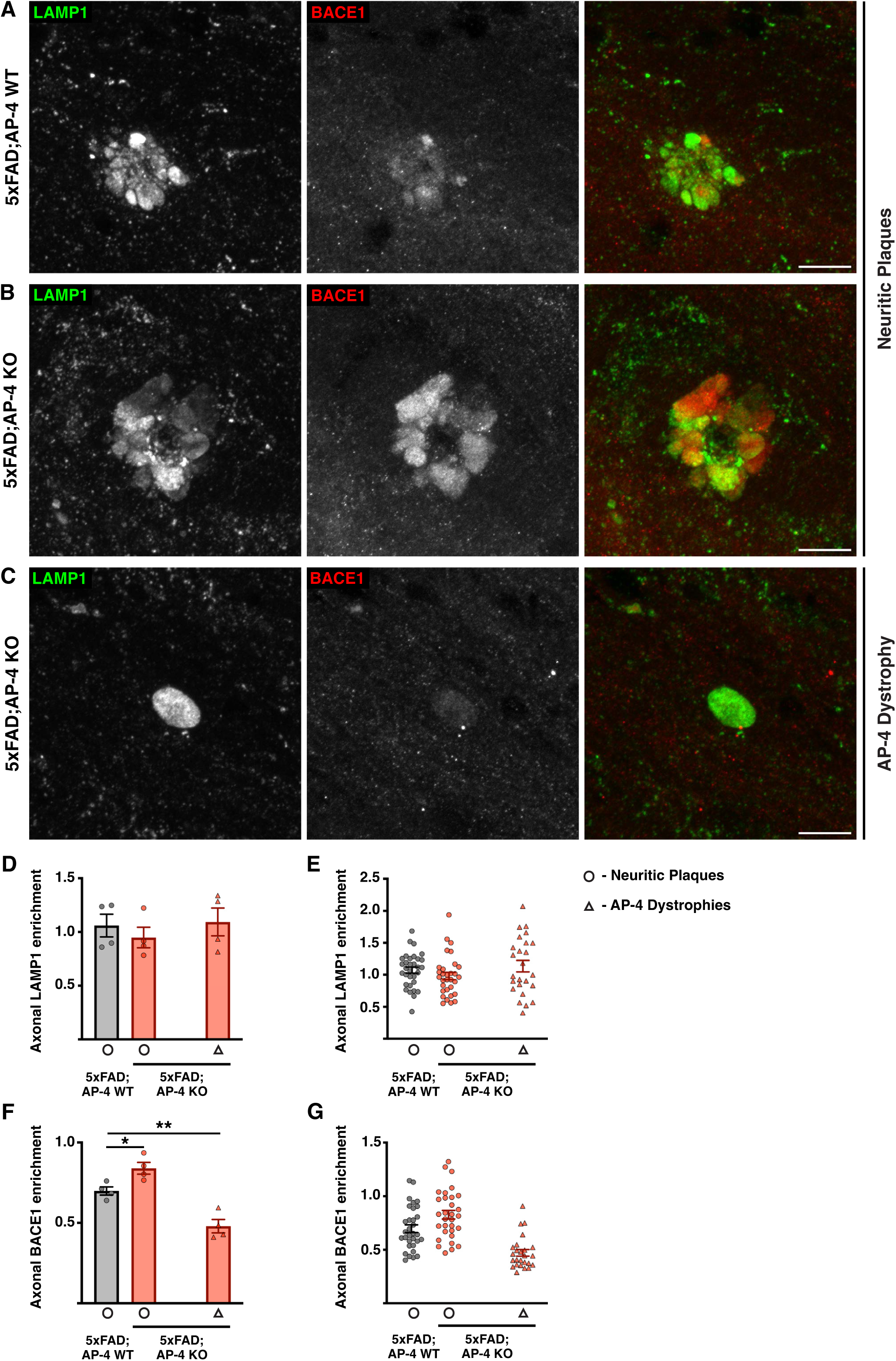
Loss of AP-4 complex increases BACE1 enrichment in amyloid plaques. (A-C) High-resolution confocal images of LAMP1 (green; lysosomes) and BACE1 (red; BACE1-positive vesicles) depicting neuritic plaques (A, B) and AP-4 dystrophies (C) in the corpus callosum of 3-month-old 5xFAD mice with AP-4 (WT; A) or lacking AP-4 (KO; B, C). Note that AP-4 dystrophies are only observed in mice lacking AP-4. Bar, 10μm. (D) Quantification of mean relative enrichment of LAMP1 in axonal dystrophies of neuritic plaques and AP-4 dystrophies compared to neuronal soma in 3-month-old female 5xFAD mice with AP-4 (WT; Grey) or lacking AP-4 (KO; Red). Mean ± SEM, N= 4, *, P < .05, one way-ANOVA with Dunnett’s post-test. Neuritic plaques-circle and AP-4 dystrophies-triangle. (E) Plot depicting relative LAMP1 enrichment from individual neuritic plaques and AP-4 dystrophies of the four animals from each genotype (Grey-5xFAD; AP-4 WT; Red-5xFAD; AP-4 KO). Neuritic plaques-circle and AP-4 dystrophies-triangle. (F) Quantification of mean relative enrichment of BACE1 in axonal dystrophies of neuritic plaques and AP-4 dystrophies compared to neuronal soma in 3-month-old female 5xFAD mice with AP-4 (WT; Grey) or lacking AP-4 (KO; Red). Mean ± SEM, N= 4, *, P < .05, one way-ANOVA with Dunnett’s post-test. Neuritic plaques-circle and AP-4 dystrophies-triangle. (G) Plot depicting relative BACE1 enrichment from individual neuritic plaques and AP-4 dystrophies of the four animals from each genotype (Grey-5xFAD; AP-4 WT; Red-5xFAD; AP-4 KO). Neuritic plaques-circle and AP-4 dystrophies-triangle.

**Figure 5:**
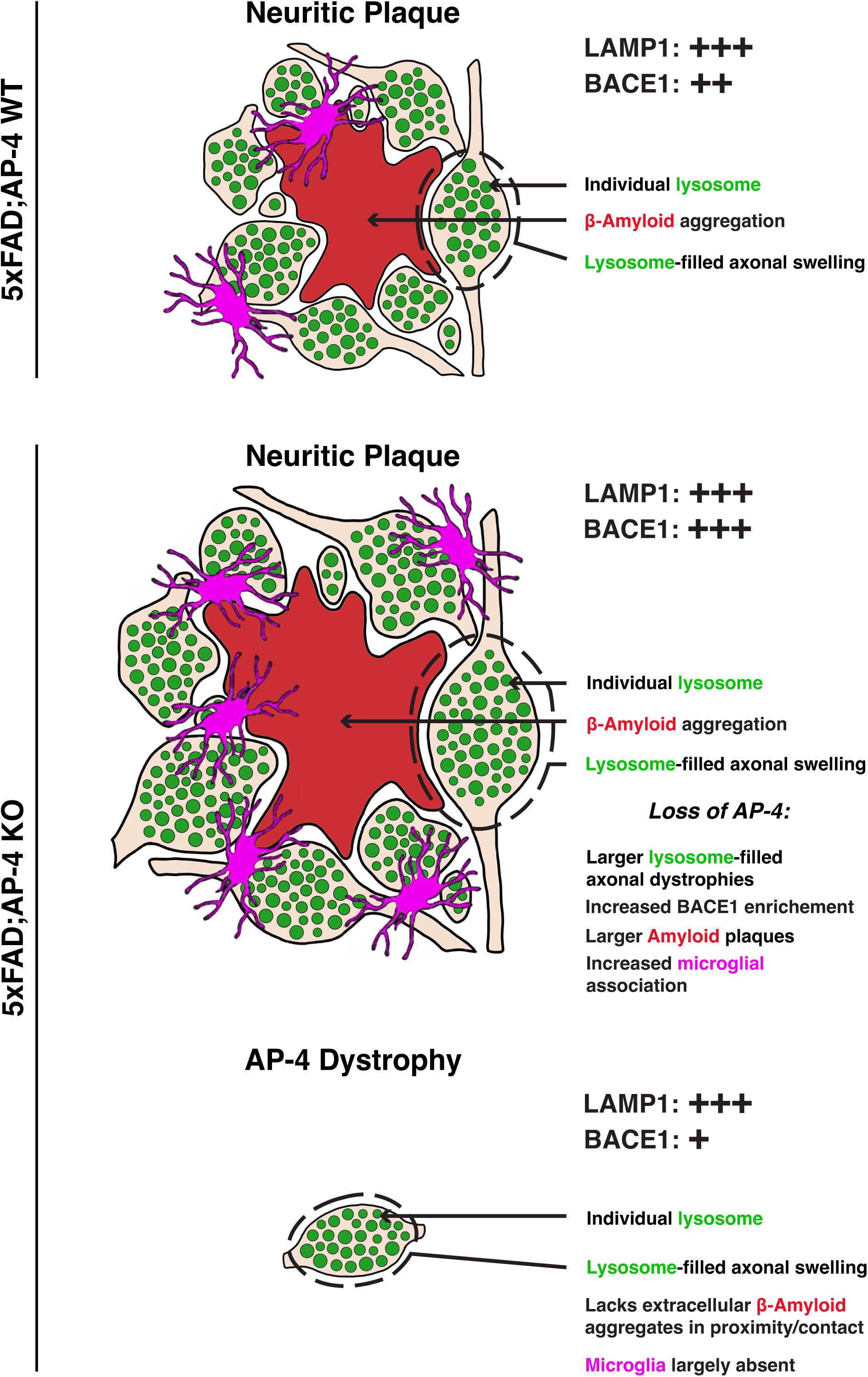
Schematic summarizing the exacerbation of neuritic plaque pathology arising from loss of AP-4 complex function in 5xFAD mice. Schematic of neuritic plaque in 5xFAD; AP-4 WT and KO animals exhibiting extracellular amyloid aggregate surrounded by lysosome-filled axonal swellings. Neuritic plaques also have microglia associated with them. AP-4 loss results in increase in size (area of both Amyloid/Aβ aggregate and lysosome-filled axonal dystrophies) of neuritic plaques as well as increases glial association with plaques. 5xFAD; AP-4 KO animals also exhibit “AP-4” dystrophies which could be seeds of future neuritic plaques. These are smaller in size, do not yet show Aβ aggregation near them, and also exhibit reduced glial association. While lysosomal protein LAMP1 is enriched in all the axonal dystrophies, BACE1 is more enriched in the neuritic plaques of 5xFAD; AP-4 KO animals and lesser in AP-4 dystrophies.

## Discussion

Our study sheds new insight into the development of neuritic plaque pathology in Alzheimer’s disease, a key pathological feature observed in both human AD and mouse models of the same (Terry et al., 1964; Nixon, 2005; Condello et al., 2011; Kandalepas et al., 2013; Gowrishankar et al., 2015). Lysosomes and lysosome-related organelles accumulating in swollen axons around amyloid plaques have been postulated to be pathologically meaningful sites of Aβ production (Gowrishankar et al., 2015; Gowrishankar et al., 2017). A recent study suggested an even more central role for lysosome dysfunction, including lysosome deacidification in neurons, in Alzheimer’s disease pathology (Lee et al., 2022). This study also proposed that the aberrant lysosomes building up around plaques arise from trapped lysosomes in membrane protrusions originating from soma of dying neurons rather than axons (Lee et al., 2022). However, another critical study that examined the role of PLD3 in the development of Alzheimer’s disease pathology, reaffirmed the axonal origins of the lysosome build up at neuritic plaques and linked their accumulation/enlargement to axonal conduction blockades (Yuan et al., 2022). These studies further highlight the importance of understanding the roles of lysosomes across the different subcellular compartments of the neuron and their disease relevance.

In recent years, the presence of axonal dystrophies containing lysosomes and their related organelles has been reported under a variety of pathological conditions, including certain forms of Hereditary Spastic Paraplegia (HSP) as well as Neiman pick type C disease (Allison et al., 2017; De Pace et al., 2018; Edmison et al., 2021; Roney et al., 2021; Majumder et al., 2022; Roney et al., 2022). The precise nature of the organelles and their composition appears to vary in some of these conditions (Roney et al., 2021; Roney et al., 2022; Paumier and Gowrishankar, 2024). Loss of the AP-4ε subunit and thus functional AP-4 complex in mice, resulted in axonal dystrophies containing LAMP1-enriched vesicles in different brain regions including the white matter tracts in the midbrain (De Pace et al., 2018; Edmison et al., 2021), which we have termed AP-4 dystrophies.

Of relevance to APP metabolism, we observed that the accumulation of axonal lysosomes was accompanied by a strong enrichment of the β secretase BACE1 as well as PSEN2 in these axonal swellings (Edmison et al., 2021) . Given the similarity in BACE1 enrichment at these AP-4 dystrophies to that observed at amyloid plaques (Zhao et al., 2007; Gowrishankar et al., 2015), and the potential for increased APP processing and Aβ production at these sites, we predicted that the axonal pathology arising from AP-4 complex loss, i.e., occurrence of AP-4 dystrophies, would lead to a worsening of the amyloid plaque pathology. Indeed, we observed that 3-month-old 5xFAD mice lacking functional AP-4 complex exhibited a dramatic worsening of amyloid plaque pathology which included an increase in size of the extracellular amyloid aggregate as well as the lysosome-filled dystrophy area (Figure 1D, F). Interestingly, this exacerbation of plaque pathology was specific to the hippocampus and corpus callosum of the animals while the pathology in the cortex was not significantly increased (Figure S3). This region-specific worsening of pathology correlated strongly with the burden/extent of axonal lysosome pathology/ AP-4 dystrophies (Figure 3G; percent area occupied by lysosome-filled axonal dystrophies), suggesting that these accumulating lysosomes could contribute to amyloidogenesis and development of amyloid pathology. Prior studies that involved genetic manipulation to deplete JIP3, a lysosome adaptor that is critical for retrograde axonal lysosome transport, in 5xFAD animals resulted in a drastic increase in amyloid plaque burden (Gowrishankar et al., 2017) . Our studies in human i^3^Neurons examining of the nature of the lysosomes and related organelles that build up in swollen axons upon AP-4 loss, led to the surprising finding of increased JIP3 in these axonal swellings (Majumder et al., 2022). While the underlying cause for JIP3 accumulation in axons remains to be determined, the massive accumulation of axonal lysosomes in this region, reminiscent of the phenotype of JIP3 KO neurons (Gowrishankar et al., 2017; Gowrishankar et al., 2021), suggests that the JIP3 building up in dystrophies of AP-4 KO neurons may not be not functioning effectively. Recent studies have identified multiple *de novo* variants in JIP3/MAPK8IP3 which are associated with a neurodevelopmental disorder presenting with mild to severe intellectual disability and other brain anomalies (Iwasawa et al., 2019; Platzer et al., 2019). Further strengthening the connection of altered axonal lysosome abundance and neuropathology, CRISPR-based editing to express some of these human mutations in *C elegans* resulted in increased axonal lysosome density in these nematodes (Platzer et al., 2019). Intriguingly, AP-4 deficiency syndrome or AP-4 associated HSP, the human disorder arising from loss of AP-4 complex function, has several overlapping clinical presentations with the JIP3-linked neurodevelopmental disorder, including intellectual disability, hypoplasia of the corpus callosum, developmental delays including problems with speech development (Ebrahimi-Fakhari et al., 1993; Iwasawa et al., 2019; Platzer et al., 2019). Despite these similarities, the composition or at the very least, the immediate environment around the accumulating axonal lysosomes under these two conditions (AP-4 complex loss versus JIP3 loss) are likely to be different with the former having high levels of JIP3 in the axonal dystrophies carrying the lysosomes while the latter completely lacks JIP3. However, despite this major difference, both conditions lead to an enhancement of amyloid pathology in 5xFAD mice and a concomitant increase in APP processing machinery including BACE1 and PSEN2 in axonal swellings. This suggests that prolonged and/or increased BACE1 presence in axonal endolysosomes could promote amyloidogenesis. We have previously shown that stalling axonal lysosomes through genetic perturbation to JIP3 does increase intraneuronal Aβ42 (Gowrishankar et al., 2017; Gowrishankar et al., 2021). In this study here, our examination of axonal enrichment of BACE1 and LAMP1 in the dystrophies at amyloid plaques revealed that the axonal organelles in AP-4 KO animals showed a stronger enrichment of BACE1 (Figure 4F, G) than control 5xFAD mice having normal AP-4 function while they had comparable levels of LAMP1 (Figure 4D, E).

AP-4 adaptor complexes also affect the sorting of different cargo from the trans-Golgi network. Of note, AP-4 has been shown to interact with exogenously expressed APP in a cell line model (Burgos et al., 2010). It remains to be established if sorting and subsequent processing of endogenous APP is altered in neurons upon AP-4 loss, and even if so, whether some neurons are more vulnerable than others. AP-4 also regulates sorting and traffic of APOER2 which is linked to Reelin signaling (Caracci et al., 2024). The effect of altered APOER2 traffic was observed in cortical and hippocampal neurons (Caracci et al., 2024). While the altered sorting of different cargo could contribute to amyloid pathology, they are likely affected in all neuronal types. Thus far, the lysosome-filled axonal dystrophy is the one pathological feature that shows strong correlation in the same region where the plaque pathology is worsened.

There were also some other striking differences between the pathologies caused by AP-4 loss when compared to JIP3 depletion. In addition to region-specific enhancement of amyloid pathology upon AP-4 loss, we also observed increased glial recruitment to plaques in 5xFAD animals lacking AP-4 (Figure 2E, I). While JIP3 haploinsufficiency caused an increase in plaque size and abundance, the glial recruitment was largely proportional to size of the plaques (Gowrishankar et al., 2017). However, loss of AP-4 caused increase glial association with neuritic plaques, with even moderately sized plaques having several microglia associated to them (Figure 2G). While it is possible that the higher frequency of plaques within the region contributes to more glia in the vicinity being attracted to the plaques, loss of the AP-4 complex could likely also alter the glia-plaque interaction. A recent study that integrated AD GWAS with myeloid epigenomic and transcriptomic data sets identified AD risk enhancers and nominated candidate causal genes that are their targets. This included genes *AP4M1* and *AP4E1*, which encode two of the subunits of the heterotetrametric complex (Novikova et al., 2021). AP-4 complex promotes autophagosome maturation in different cell types through export of critical autophagy protein (Mattera et al., 2017; Davies et al., 2018) ATG9. As microglial autophagy is activated in mouse models of AD, alteration to this pathway due to AP-4 loss could affect microglia-plaque interactions (Choi et al., 2023). However, inhibition of microglial autophagy causes disengagement of microglia from plaques (Choi et al., 2023) which is distinct from the phenotype observed upon AP-4 loss (Figure 2F-I). Intriguingly, acute treatment with aducanumab in APP/PS1 mice resulted in decrease in amyloid and LAMP1-associated neuritic dystrophy but was accompanied by increase microglial association with plaques, suggesting a more complex relationship between glial recruitment and plaque development and clearance (Cadiz et al., 2024). Future studies involving microglia-specific knock out of AP-4ε will help discern the contribution from glia to the development of amyloid pathology arising from AP-4 loss. However, in support of a more central contribution from the axonal lysosome pathology, there is a strong correlation between areas exhibiting increased AP-4 dystrophy-burden in AP-4ε KO animals with where the amyloid plaque pathology is worsened in 5xFAD; AP-4 KO animals; whereas gliosis is not significantly enhanced in AP-4ε KO animals (Edmison et al., 2021). The AP-4 dystrophies, persist in the 5xFAD; AP-4 KO animals as well, and are distinguishable from the neuritic plaques in being smaller, lacking aggregated Aβ in their immediate vicinity and largely being devoid of glial association (Figure 3). Given their relatively lower enrichment of BACE1 compared to dystrophic axons of neuritic plaques in either genotype, these would likely have lower levels of Aβ production. This, along with their comparatively sparser distribution may not favor aggregate formation at least in 3-month-old animals. Interestingly, AP-4 dystrophies have a higher enrichment of the lysosomal GTPase, Arl8, when compared to the dystrophies in neuritic plaques of either genotype (Figure S9). The significance of the enrichment of this key lysosomal GTPase that is linked to both kinesin-and dynein-dependent lysosome transport as well multiple fusion events at the lysosome (Khatter et al., 2015; Ballabio and Bonifacino, 2020), needs to be determined and is the focus of future studies. Of relevance to human Alzheimer’s disease, unbiased proteomics carried out on amyloid plaques from Early onset Alzheimer’s disease (EOAD) patients identified Arl8 as a protein that is highly enriched at plaques (Drummond et al., 2022). A second study that examined proteins changing in human AD as well as mouse models of AD also identified Arl8 as a highly enriched protein at amyloid plaques (Boeddrich et al., 2023). While the study suggested that the Arl8 enrichment could be a compensatory response to lysosome accumulation, the activity status of Arl8, including membrane recruitment, interaction with effectors, will help determine the functional consequences of higher Arl8 levels in these dystrophies.

Our study, in conjunction with others, supports a model wherein axonal lysosomes contribute to Aβ production and thus AD pathogenesis. Recently, a high throughput screen identified a small molecule that rescued ATG9 traffic in AP-4 deficient cells (Saffari et al., 2024). This correction of ATG9 traffic was observed in patient fibroblasts as well as iPSC-derived neurons (Saffari et al., 2024). Examining the effect of the small molecule on BACE1 and LAMP1 accumulation in dystrophic axons as well as on amyloid plaque pathology could shed more light on the contribution of these pathways to AD pathogenesis and the therapeutic potential of manipulating AP-4-dependent pathways in AD treatment.

## Methods

### Mouse Strains

Animal procedures were approved by and carried out in accordance with guidelines established by the Office of Animal Care and Institutional Biosafety (OACIB), UIC. The 5xFAD mice (Oakley et al., 2006) was purchased from Jackson laboratory, and C57BL/6N-Ap4ε1tm1b(KOMP)Wtsi was purchased from the European Mouse Mutant Archive (EMMA) mice have been described previously (Edmison et al., 2021; Davies et al., 2022; Park et al., 2023). Male heterozygous 5xFAD mice were bred with heterozygous AP-4 females, to generate 5xFAD/+; AP4 +/-males which were in turn mated with heterozygous AP-4 females to generate 5xFAD; AP-4 WT, 5xFAD; AP-4 KO mice (progenies genotyped by PCR). AP-4 genotyping was performed in-house with the following primers: AP4E1-5arm-WTF(5’GCCTCTGTTTAGTTTGCGATG3’), AP4E1-Crit-WTR (5’CGTGCACAGACAGGTTTGAT3’), 5mut-R1(5’GAACTTCGGAATAGGAACTTCG3’). 5xFAD progenies were genotyped via Transnetyx with the following probes: APPsw Tg and huPSEN1 Tg. These animals were then aged to 3 months and sex-matched animals of the two genotypes were used for experiments.

### Transcardial Perfusion

Transcardial perfusion was performed to isolate tissue for immunohistochemistry experiments. PBS was administered to anesthetized mice (Isoflurane) by transcardial perfusion. The isolated brains were then quickly dissected sagittally down the midline to procure hemi-brains, and one half fixed by immersion in 4% paraformaldehyde (PFA) overnight at 4°C while the other half was flash frozen in liquid nitrogen and stored at -80°C for future biochemical studies.

### Immunohistochemistry Studies

Hemibrains that were fixed in PFA overnight were washed 3x times in PBS and sectioned coronally at 30μm thickness using a vibratome (Leica VT1200S). Sections were collected serially into 8 wells of a 24-well plate containing PBS containing .05% sodium azide. Sections of the mid brain (sections corresponding to coronal sections 69-74 in the Allen mouse brain atlas) were selected from the appropriate well and used for staining after matching them according to their depth. Staining experiments on the free-floating sections were performed as described previously (Gowrishankar et al., 2015).

### Microscopy

Stitched images of immunostained brain sections were acquired using the BZ-X800, Keyence microscope equipped with a 20x, .75 NA objective or using a laser scanning confocal microscope (LSM 880; ZEISS) equipped with a 63X plan Apo (NA 1.4) oil immersion objective. High resolution images of individual plaques and glia were acquired using a laser scanning confocal microscope (LSM 710; ZEISS equipped with a 63X plan Apo (NA 1.4) oil immersion objective lens or BZ-X800, Keyence microscope equipped with a 60x BZ Series infinite optical system oil-immersion objective lens. Z-stacks with a step size of 0.2 and 0.1 μm were routinely acquired with the confocal microscope and BZ-X800, respectively.

### Quantification of Axonal Lysosome Accumulation and Plaque Area

Coronal brain sections from 3-month-old 5xFAD; AP-4 WT and 5xFAD; AP-4 KO mice were co-stained for LAMP1 and Aβ and imaged using BZ-X800, Keyence microscope. Images (Z-stack images that encompassed LAMP1 and Aβ signals, typically 30-50 planes with 0.1-μm step size) of every individual amyloid plaque and the LAMP1 accumulations around them were acquired (60× objective) from the corpus callosum and hippocampus. Images were processed using BZ-X800 Analyzer software which takes each image from the Z-stack and combines the highest contrast portions of each slice to generate a fully focused 2D image. The images of the LAMP1 and amyloid staining were analyzed using ImageJ software, where regions of interest were manually outlined, and the area associated with the outlined region was computed.

### Quantification of Microglial Association with Amyloid Plaques

Coronal sections from 3-month-old 5xFAD; AP-4 WT and 5xFAD; AP-4 KO mice were costained for LAMP1 (lysosomes) and iba1 (microglia), and imaged by BZ-X800, Keyence microscope using a 60× objective (NA 1.4). Images (Z-stack images that encompassed LAMP1 and iba1 signals, typically 30-50 planes with 0.1-μm step size) of all plaques from the corpus callosum and hippocampus were acquired. Images of the two channels were overlaid and analyzed using ImageJ software where regions of interest (LAMP1 signal) were manually outlined, and the area associated with the outlined region was computed to determine size of plaques. All microglia with any overlap of iba1 signal with LAMP1 of a plaque were scored as associated with that specific plaque.

### Measurement of Lysosome Protein Enrichment at Axonal Dystrophies

Lysosomal protein enrichment in axonal dystrophies was measured as described previously (Gowrishankar et al., 2015). Briefly, brain sections from 3-month-old 5xFAD; AP-4 WT and 5xFAD; AP-4 KO mice were costained for LAMP1 and BACE1 and high-resolution images were acquired by laser-scanning confocal microscopy (63× objective). Z-stacks were acquired to encompass a selected amyloid plaque (typically 12–15 Z-planes with a step size of 0.419 μm). ImageJ software was used to calculate the enrichment of the respective lysosomal proteins in the dystrophies. To this end, 20-30 regions of interest were outlined within each amyloid plaque axonal dystrophies and AP-4 dystrophies and over cell body lysosomes from each animal and between seven to ten plaques/AP-4 dystrophies and five to six soma per animal per genotype was examined in this way. The mean intensity in each such region was determined, and the ratio between them was calculated for each of these proteins. Four different mice were analyzed for each genotype.

### Tissue Expansion and Imaging

The stained brain sections were incubated in the anchoring reagent Acryloyl-X, SE (AcX) (A20770, Thermo Fisher) (0.1 mg/mL AcX in 1x PBS) overnight at room temperature (RT) (Asano et al., 2018).To remove unreacted AcX, the brain sections were washed twice with 1x PBS for 15 min per wash. For gelation, the brain sections were incubated in gelling solution [1x PBS, 2 M NaCl, 8.625% (w/v) sodium acrylate, 2.5% (w/v) acrylamide, 0.15% (w/v) N,N′-methylenebisacrylamide, 0.01% (w/v) of 4-hydroxy-2,2,6,6-tetramethylpiperidin-1-oxyl (4HT), 0.2% (w/v) of ammonium persulfate (APS), and 0.2% (w/v) of tetramethylethylenediamine (TEMED)] at 4°C for 20 min. The sections were then allowed to gel in a humidified 37°C incubator for 1-1.5 hours. Immediately, the gelled sections were immersed in Proteinase K digestion buffer (1:100 dilution to a final concentration of 800U = 8 U/ml) overnight at RT for homogenization^1^. The gelled sections were then washed three times with water, 20 min per wash, for expansion. Images of the expanded brain sections were captured using a Nikon spinning disk confocal microscope (CSU-W1, Yokogawa) with 40x, 1.15 NA water-immersion objective (Nikon).

### Statistical Methods

Data represent mean ± SEM unless otherwise specified. All statistical tests were performed using GraphPad Prism 10 software. Statistical tests performed are indicated in their respective Figure legends [including the number of animals (N), statistical test used, and p values].

## Acknowledgements

Research reported in this publication was supported by the Wolverine Foundation (Swetha Gowrishankar/ S.G), The Ralph and Marian Falk Medical Trust (S.G), the National Institute on Aging of the National Institutes of Health [RF1AG076653 (S.G), R01AG074248 (S.G)], National Institutes of Health [DP2MH136390 (Ruixuan Gao/ R.G), UG3MH126864 (R.G)], Searle Scholars Program (R.G), McKnight Technological Innovations in Neuroscience Award (R.G). The content is solely the responsibility of the authors and does not necessarily represent the official views of the National Institutes of Health.

## Conflict of Interest Statement

R.G. is a co-inventor of multiple patents related to expansion microscopy. The other authors declare that they have no competing financial interests.

## Notes

### Summary of Updates

The revised manuscript includes further characerization of the AP-4 dystrophies. We show that they share similarities with the FAD dystrophies in terms of luminal proteases, lysosomal GTPase RagC. Interestingly, we also find high levels of the small GTPase, Arl8, enriched in AP-4 dystrophies.

## References

Allison R, Edgar JR, Pearson G, Rizo T, Newton T, Gunther S, Berner F, Hague J, Connell JW, Winkler J, Lippincott-Schwartz J, Beetz C, Winner B, Reid E (2017) Defects in ER-endosome contacts impact lysosome function in hereditary spastic paraplegia. J Cell Biol 216:1337–1355.

Asano SM, Gao R, Wassie AT, Tillberg PW, Chen F, Boyden ES (2018) Expansion Microscopy: Protocols for Imaging Proteins and RNA in Cells and Tissues. Curr Protoc Cell Biol 80:e56.

Ballabio A, Bonifacino JS (2020) Lysosomes as dynamic regulators of cell and organismal homeostasis. Nat Rev Mol Cell Biol 21:101–118.

Boeddrich A et al. (2023) A proteomics analysis of 5xFAD mouse brain regions reveals the lysosome-associated protein Arl8b as a candidate biomarker for Alzheimer’s disease. Genome Med 15:50.

Burgos PV, Mardones GA, Rojas AL, daSilva LL, Prabhu Y, Hurley JH, Bonifacino JS (2010) Sorting of the Alzheimer’s disease amyloid precursor protein mediated by the AP-4 complex. Dev Cell 18:425–436.

Cadiz MP, Gibson KA, Todd KT, Nascari DG, Massa N, Lilley MT, Olney KC, Al-Amin MM, Jiang H, Holtzman DM, Fryer JD (2024) Aducanumab anti-amyloid immunotherapy induces sustained microglial and immune alterations. J Exp Med 221.

Caracci MO, Pizarro H, Alarcon-Godoy C, Fuentealba LM, Farfan P, De Pace R, Santibanez N, Cavieres VA, Pastor TP, Bonifacino JS, Mardones GA, Marzolo MP (2024) The Reelin receptor ApoER2 is a cargo for the adaptor protein complex AP-4: Implications for Hereditary Spastic Paraplegia. Prog Neurobiol 234:102575.

Cataldo AM, Hamilton DJ, Nixon RA (1994) Lysosomal abnormalities in degenerating neurons link neuronal compromise to senile plaque development in Alzheimer disease. Brain Res 640:68–80.

Choi I, Wang M, Yoo S, Xu P, Seegobin SP, Li X, Han X, Wang Q, Peng J, Zhang B, Yue Z (2023) Autophagy enables microglia to engage amyloid plaques and prevents microglial senescence. Nat Cell Biol 25:963–974.

Condello C, Schain A, Grutzendler J (2011) Multicolor time-stamp reveals the dynamics and toxicity of amyloid deposition. Sci Rep 1:19.

Cras P, Kawai M, Lowery D, Gonzalez-DeWhitt P, Greenberg B, Perry G (1991) Senile plaque neurites in Alzheimer disease accumulate amyloid precursor protein. Proc Natl Acad Sci U S A 88:7552–7556.

Cummings BJ, Su JH, Geddes JW, Van Nostrand WE, Wagner SL, Cunningham DD, Cotman CW (1992) Aggregation of the amyloid precursor protein within degenerating neurons and dystrophic neurites in Alzheimer’s disease. Neuroscience 48:763–777.

Davies AK, Itzhak DN, Edgar JR, Archuleta TL, Hirst J, Jackson LP, Robinson MS, Borner GHH (2018) AP-4 vesicles contribute to spatial control of autophagy via RUSC-dependent peripheral delivery of ATG9A. Nat Commun 9:3958.

Davies AK, Alecu JE, Ziegler M, Vasilopoulou CG, Merciai F, Jumo H, Afshar-Saber W, Sahin M, Ebrahimi-Fakhari D, Borner GHH (2022) AP-4-mediated axonal transport controls endocannabinoid production in neurons. Nat Commun 13:1058.

De Pace R, Skirzewski M, Damme M, Mattera R, Mercurio J, Foster AM, Cuitino L, Jarnik M, Hoffmann V, Morris HD, Han TU, Mancini GMS, Buonanno A, Bonifacino JS (2018) Altered distribution of ATG9A and accumulation of axonal aggregates in neurons from a mouse model of AP-4 deficiency syndrome. PLoS Genet 14:e1007363.

Dell’Angelica EC, Mullins C, Bonifacino JS (1999) AP-4, a novel protein complex related to clathrin adaptors. J Biol Chem 274:7278–7285.

Drerup CM, Nechiporuk AV (2013) JNK-interacting protein 3 mediates the retrograde transport of activated c-Jun N-terminal kinase and lysosomes. PLoS Genet 9:e1003303.

Drummond E, Kavanagh T, Pires G, Marta-Ariza M, Kanshin E, Nayak S, Faustin A, Berdah V, Ueberheide B, Wisniewski T (2022) The amyloid plaque proteome in early onset Alzheimer’s disease and Down syndrome. Acta Neuropathol Commun 10:53.

Ebrahimi-Fakhari D, Behne R, Davies AK, Hirst J (1993) AP-4-Associated Hereditary Spastic Paraplegia. In: GeneReviews((R)) (Adam MP, Feldman J, Mirzaa GM, Pagon RA, Wallace SE, Bean LJH, Gripp KW, Amemiya A, eds). Seattle (WA).

Edmison D, Wang L, Gowrishankar S (2021) Lysosome Function and Dysfunction in Hereditary Spastic Paraplegias. Brain Sci 11.

Edwards SL, Yu SC, Hoover CM, Phillips BC, Richmond JE, Miller KG (2013) An Organelle Gatekeeper Function for Caenorhabditis elegans UNC-16 (JIP3) at the Axon Initial Segment. Genetics 194:143–161.

Fiala JC (2007) Mechanisms of amyloid plaque pathogenesis. Acta Neuropathol 114:551–571.

Gowrishankar S, Wu Y, Ferguson SM (2017) Impaired JIP3-dependent axonal lysosome transport promotes amyloid plaque pathology. J Cell Biol.

Gowrishankar S, Lyons L, Rafiq NM, Roczniak-Ferguson A, De Camilli P, Ferguson SM (2021) Overlapping roles of JIP3 and JIP4 in promoting axonal transport of lysosomes in human iPSC-derived neurons. Mol Biol Cell:mbcE20060382.

Gowrishankar S, Yuan P, Wu Y, Schrag M, Paradise S, Grutzendler J, De Camilli P, Ferguson SM (2015) Massive accumulation of luminal protease-deficient axonal lysosomes at Alzheimer’s disease amyloid plaques. Proc Natl Acad Sci U S A 112:E3699–3708.

Hirst J, Bright NA, Rous B, Robinson MS (1999) Characterization of a fourth adaptor-related protein complex. Mol Biol Cell 10:2787–2802.

Itagaki S, McGeer PL, Akiyama H, Zhu S, Selkoe D (1989) Relationship of microglia and astrocytes to amyloid deposits of Alzheimer disease. J Neuroimmunol 24:173–182.

Ivankovic D, Drew J, Lesept F, White IJ, Lopez Domenech G, Tooze SA, Kittler JT (2020) Axonal autophagosome maturation defect through failure of ATG9A sorting underpins pathology in AP-4 deficiency syndrome. Autophagy 16:391–407.

Iwasawa S et al. (2019) Recurrent de novo MAPK8IP3 variants cause neurological phenotypes. Ann Neurol 85:927–933.

Kandalepas PC, Sadleir KR, Eimer WA, Zhao J, Nicholson DA, Vassar R (2013) The Alzheimer’s beta-secretase BACE1 localizes to normal presynaptic terminals and to dystrophic presynaptic terminals surrounding amyloid plaques. Acta Neuropathol 126:329–352.

Khatter D, Sindhwani A, Sharma M (2015) Arf-like GTPase Arl8: Moving from the periphery to the center of lysosomal biology. Cell Logist 5:e1086501.

Lee JH, Yang DS, Goulbourne CN, Im E, Stavrides P, Pensalfini A, Chan H, Bouchet-Marquis C, Bleiwas C, Berg MJ, Huo C, Peddy J, Pawlik M, Levy E, Rao M, Staufenbiel M, Nixon RA (2022) Faulty autolysosome acidification in Alzheimer’s disease mouse models induces autophagic build-up of Abeta in neurons, yielding senile plaques. Nat Neurosci 25:688–701.

Leng F, Edison P (2021) Neuroinflammation and microglial activation in Alzheimer disease: where do we go from here? Nat Rev Neurol 17:157–172.

Majumder P, Edmison D, Rodger C, Patel S, Reid E, Gowrishankar S (2022) AP-4 regulates neuronal lysosome composition, function, and transport via regulating export of critical lysosome receptor proteins at the trans-Golgi network. Mol Biol Cell 33:ar102.

Mattera R, Park SY, De Pace R, Guardia CM, Bonifacino JS (2017) AP-4 mediates export of ATG9A from the trans-Golgi network to promote autophagosome formation. Proc Natl Acad Sci U S A 114:E10697–E10706.

Nixon RA (2005) Endosome function and dysfunction in Alzheimer’s disease and other neurodegenerative diseases. Neurobiol Aging 26:373–382.

Novikova G, Kapoor M, Tcw J, Abud EM, Efthymiou AG, Chen SX, Cheng H, Fullard JF, Bendl J, Liu Y, Roussos P, Bjorkegren JL, Liu Y, Poon WW, Hao K, Marcora E, Goate AM (2021) Integration of Alzheimer’s disease genetics and myeloid genomics identifies disease risk regulatory elements and genes. Nat Commun 12:1610.

Oakley H, Cole SL, Logan S, Maus E, Shao P, Craft J, Guillozet-Bongaarts A, Ohno M, Disterhoft J, Van Eldik L, Berry R, Vassar R (2006) Intraneuronal beta-amyloid aggregates, neurodegeneration, and neuron loss in transgenic mice with five familial Alzheimer’s disease mutations: potential factors in amyloid plaque formation. J Neurosci 26:10129–10140.

Park D, Wu Y, Wang X, Gowrishankar S, Baublis A, De Camilli P (2023) Synaptic vesicle proteins and ATG9A self-organize in distinct vesicle phases within synapsin condensates. Nat Commun 14:455.

Paumier JM, Gowrishankar S (2024) Disruptions in axonal lysosome transport and its contribution to neurological disease. Curr Opin Cell Biol 89:102382.

Platzer K et al. (2019) De Novo Variants in MAPK8IP3 Cause Intellectual Disability with Variable Brain Anomalies. Am J Hum Genet 104:203–212.

Roney JC, Cheng XT, Sheng ZH (2022) Neuronal endolysosomal transport and lysosomal functionality in maintaining axonostasis. J Cell Biol 221.

Roney JC, Li S, Farfel-Becker T, Huang N, Sun T, Xie Y, Cheng XT, Lin MY, Platt FM, Sheng ZH (2021) Lipid-mediated motor-adaptor sequestration impairs axonal lysosome delivery leading to autophagic stress and dystrophy in Niemann-Pick type C. Dev Cell 56:1452–1468 e1458.

Saffari A, Brechmann B, Boger C, Saber WA, Jumo H, Whye D, Wood D, Wahlster L, Alecu JE, Ziegler M, Scheffold M, Winden K, Hubbs J, Buttermore ED, Barrett L, Borner GHH, Davies AK, Ebrahimi-Fakhari D, Sahin M (2024) High-content screening identifies a small molecule that restores AP-4-dependent protein trafficking in neuronal models of AP-4-associated hereditary spastic paraplegia. Nat Commun 15:584.

Sannerud R et al. (2016) Restricted Location of PSEN2/gamma-Secretase Determines Substrate Specificity and Generates an Intracellular Abeta Pool. Cell 166:193–208.

Sharoar MG, Hu X, Ma XM, Zhu X, Yan R (2019) Sequential formation of different layers of dystrophic neurites in Alzheimer’s brains. Mol Psychiatry 24:1369–1382.

Song WM, Colonna M (2018) The identity and function of microglia in neurodegeneration. Nat Immunol 19:1048–1058.

Su JH, Cummings BJ, Cotman CW (1993) Identification and distribution of axonal dystrophic neurites in Alzheimer’s disease. Brain Res 625:228–237.

Terry RD, Gonatas NK, Weiss M (1964) Ultrastructural Studies in Alzheimer’s Presenile Dementia. Am J Pathol 44:269–297.

Yoon SY, Choi JU, Cho MH, Yang KM, Ha H, Chung IJ, Cho GS, Kim DH (2013) alpha-secretase cleaved amyloid precursor protein (APP) accumulates in cholinergic dystrophic neurites in normal, aged hippocampus. Neuropathol Appl Neurobiol 39:800–816.

Yu WH, Cuervo AM, Kumar A, Peterhoff CM, Schmidt SD, Lee JH, Mohan PS, Mercken M, Farmery MR, Tjernberg LO, Jiang Y, Duff K, Uchiyama Y, Naslund J, Mathews PM, Cataldo AM, Nixon RA (2005) Macroautophagy--a novel Beta-amyloid peptide-generating pathway activated in Alzheimer’s disease. J Cell Biol 171:87–98.

Yuan P, Zhang M, Tong L, Morse TM, McDougal RA, Ding H, Chan D, Cai Y, Grutzendler J (2022) PLD3 affects axonal spheroids and network defects in Alzheimer’s disease. Nature 612:328–337.

Zhao J, Fu Y, Yasvoina M, Shao P, Hitt B, O’Connor T, Logan S, Maus E, Citron M, Berry R, Binder L, Vassar R (2007) Beta-site amyloid precursor protein cleaving enzyme 1 levels become elevated in neurons around amyloid plaques: implications for Alzheimer’s disease pathogenesis. J Neurosci 27:3639–3649.

