## Supplemental Figures and Legends for "Axonal organelle buildup from loss of AP-4 complex function causes exacerbation of amyloid plaque pathology and gliosis in Alzheimer’s disease mouse model"

**Table S1. Antibody Summary**

| Antibody | Source | Catalog number | Dilution |
|---|---|---|---|
| Amyloid-β | Cell Signaling Technology | 2454 | 1:500 |
| ARL8B | Invitrogen | PA5-98885 | 1:200 |
| BACE1 | Cell Signaling Technology | D10E5 | 1:100 |
| Cathepsin B | R&D systems | AF965 | 1:400 |
| Iba-1 | Wako | 019-19741 | 1:250 |
| LAMP1 | DSHB | 1D4B | 1:500 |
| mLegumain | R&D systems | AF2058 | 1:100 |
| mPGRN | R&D systems | AF2557 | 1:200 |
| RagC | Cell Signaling Technology | 9480S | 1:200 |

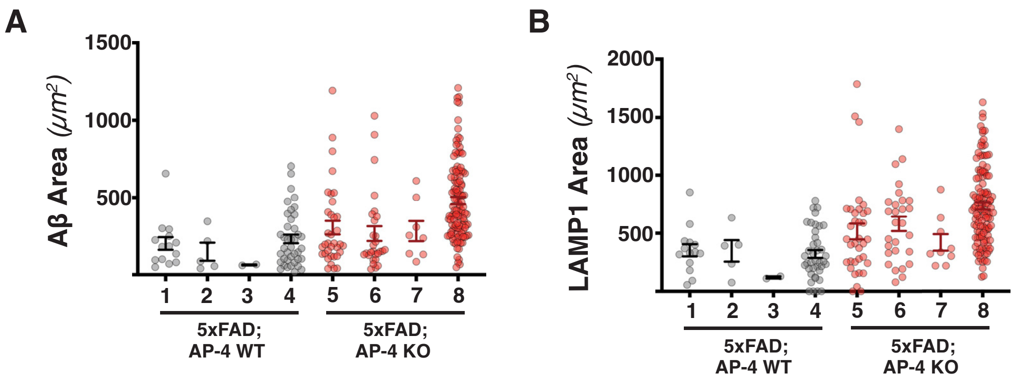

A
1500
1000
500
Aβ Area (μm²)
1
2
3
4
5
6
7
8
5xFAD;
AP-4 WT
5xFAD;
AP-4 KO
B
2000
1500
1000
500
LAMP1 Area (μm²)
1
2
3
4
5
6
7
8
5xFAD;
AP-4 WT
5xFAD;
AP-4 KO


**Figure S1**

**Figure S1**: **Loss of AP-4 complex increases size of amyloid plaques in individual 3-month-old female 5xFAD mice**

(A) Plot of areas of every individual amyloid aggregate present in the hippocampus and corpus callosum from a representative section of each of the four animals from each genotype (Grey- 5xFAD; AP-4 WT; Red- 5xFAD; AP-4 KO) of 3-month-old female mice. (B) Plot of the areas of individual lysosome-filled axonal swelling per each animal obtained from representative sections of the four animals from each genotype (Grey- 5xFAD; AP-4 WT; Red- 5xFAD; AP-4 KO) of 3-month-old female mice.

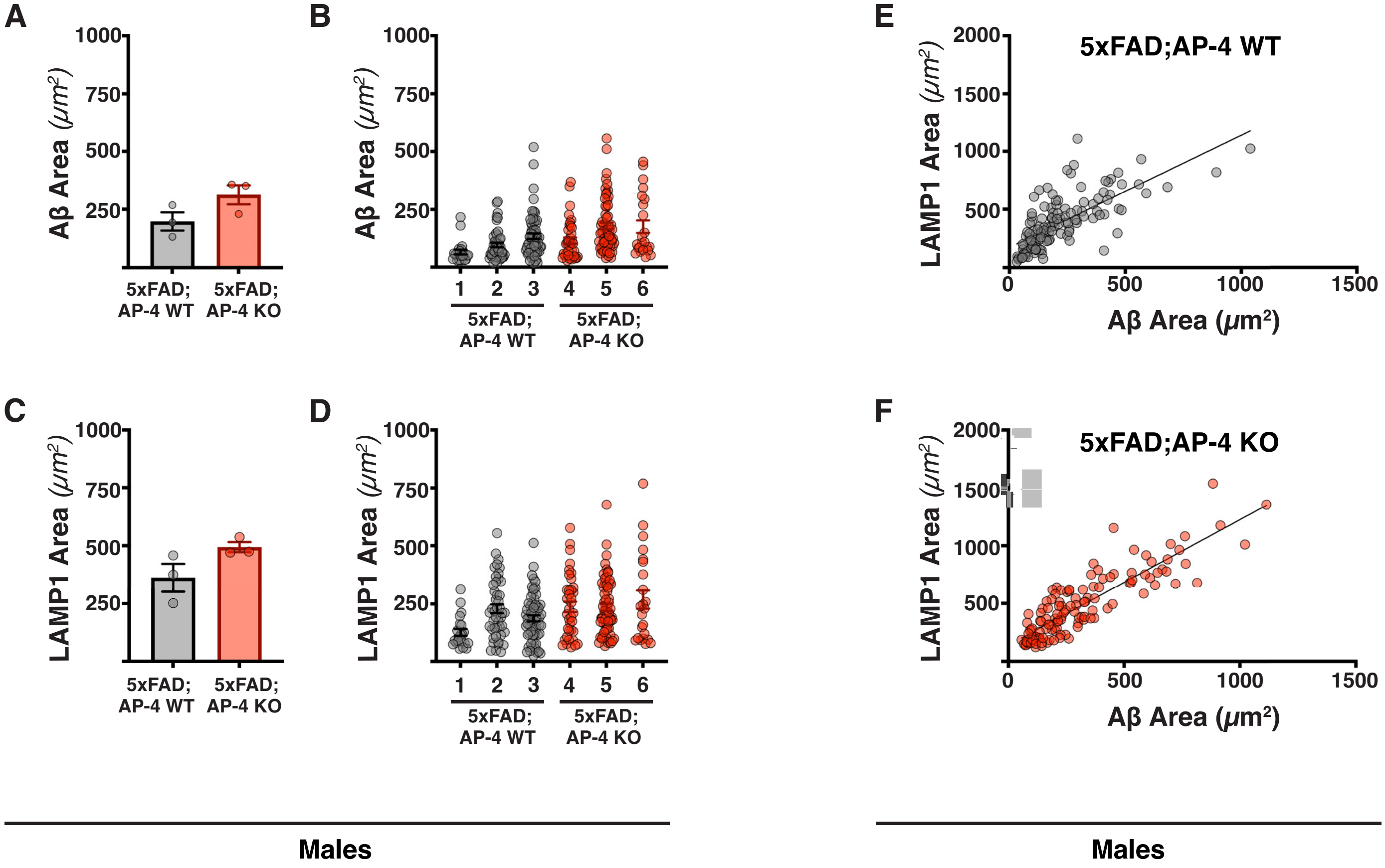

A
B
C
D
E
F
Aβ Area (μm²)
LAMP1 Area (μm²)
1000
750
500
250
2000
1500
0
5xFAD; AP-4 WT
5xFAD; AP-4 KO
1 2 3 4 5 6
5xFAD;AP-4 WT
5xFAD;AP-4 KO
Males
Males


Figure S2

**Figure S2: Loss of AP-4 complex leads to a trend towards increased size of amyloid plaques in 3-month-old male 5xFAD mice**

(A) Quantification of mean size of amyloid aggregates of neuritic plaques in 3-mo-old male 5xFAD; AP-4 WT and KO mice (Grey- AP-4 WT; Red- AP-4 KO). Mean ± SEM, N = 3 pairs of 5xFAD; AP-4 WT and 5xFAD; AP-4 KO animals. (B) Plot of areas of each individual amyloid aggregate per each animal present in the hippocampus and corpus callosum of the three animals from each genotype. (C) Quantification of mean size of lysosome-filled axonal swellings of neuritic plaques in 3-mo-old male 5xFAD; AP-4 WT and KO mice (Grey- AP-4 WT; Red- AP-4 KO). Mean ± SEM, N = 3 pairs of 5xFAD; AP-4 WT and 5xFAD; AP-4 KO animals. (D) Plot of the areas of individual lysosome-filled axonal swellings obtained from representative sections of the three animals from each genotype (Grey- 5xFAD; AP-4 WT; Red- 5xFAD; AP-4 KO). (E and F) Plot depicting the correlation between LAMP1 and Aβ areas of all individual neuritic plaques present in the hippocampus and corpus callosum of the three animals from each genotype-5xFAD; AP-4 WT (E) and 5xFAD; AP-4 KO animals (F).

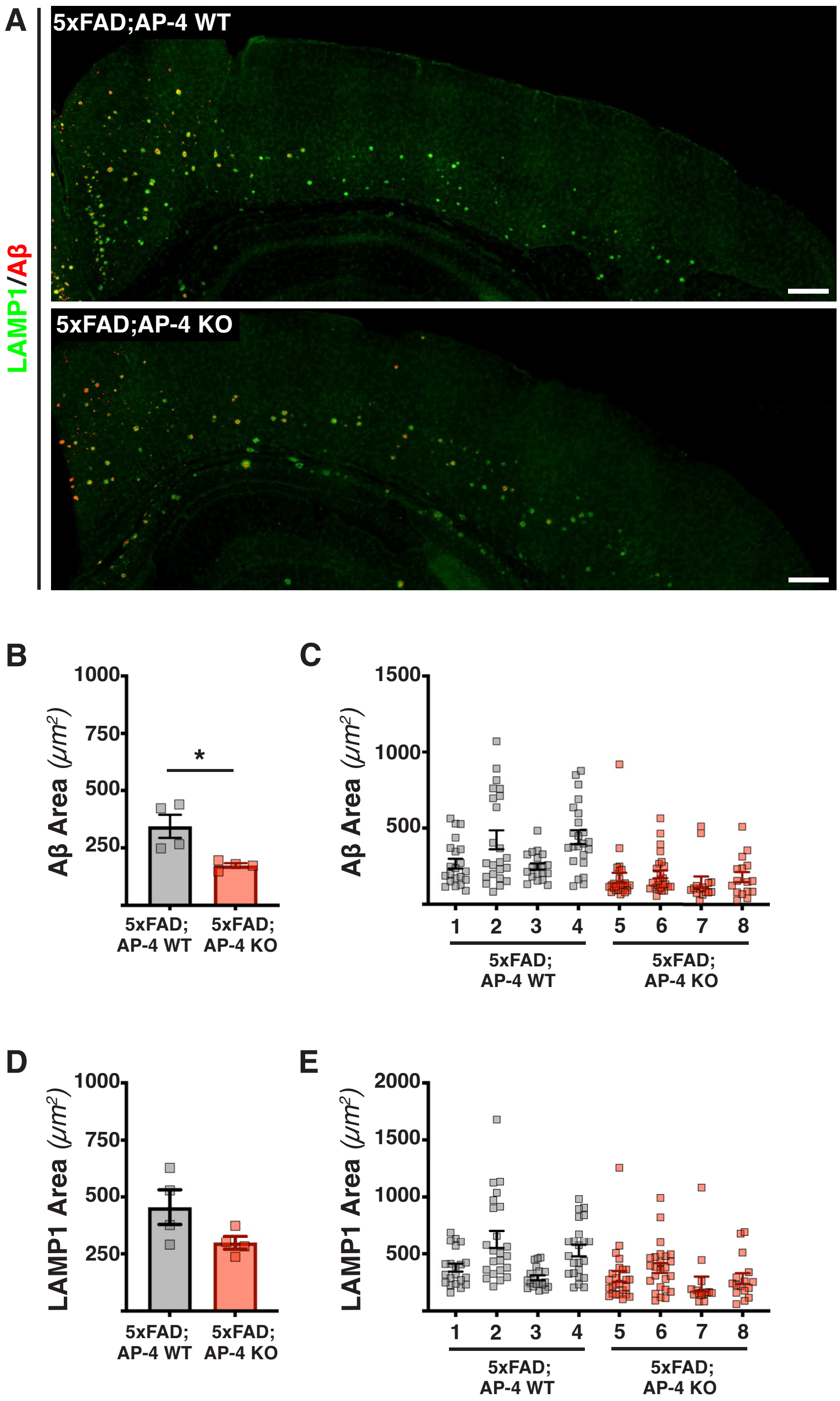
A
5xFAD;AP-4 WT
5xFAD;AP-4 KO
LAMP1/Aβ
B
Aβ Area (μm2)
1000
750
500
250
*
5xFAD; AP-4 WT
5xFAD; AP-4 KO
C
1500
1000
500
1 2 3 4 5 6 7 8
D
LAMP1 Area (μm2)
E
2000
1500

**Figure S3**

**Figure S3: Loss of AP-4 complex does not cause an exacerbation of amyloid plaque pathology in the cortex of 3-month-old 5xFAD mice**

(A) Stitched images of the cortex of 3-month-old female 5xFAD mice with AP-4 (WT) or lacking AP-4 (KO) stained for LAMP1 (green; lysosomes) and Aβ (red; amyloid aggregates) depicting neuritic plaques. Bar, 100μm. (B) Quantification of mean size of amyloid aggregates of neuritic plaques in the cortex of 3-mo-old female 5xFAD; AP-4 WT and KO mice (Grey- AP-4 WT; Red- AP-4 KO). Mean ± SEM, N = 4 pairs of 5xFAD; AP-4 WT and 5xFAD; AP-4 KO animals; *, $P < .05$, unpaired *t* test. (C) Plot of areas of each individual amyloid aggregate per each animal present in the cortex of the four animals from each genotype. (D) Quantification of mean size of lysosome-filled axonal swellings of neuritic plaques in the cortex of 3-mo-old female 5xFAD; AP-4 WT and KO mice (Grey- 5xFAD; AP-4 WT; Red- 5xFAD; AP-4 KO). Mean ± SEM, N = 4 pairs of 5xFAD; AP-4 WT and 5xFAD; AP-4 KO animals. (E) Plot of the areas of individual lysosome-filled axonal swellings of neuritic plaques per each animal present in the cortex of the four animals from each genotype.

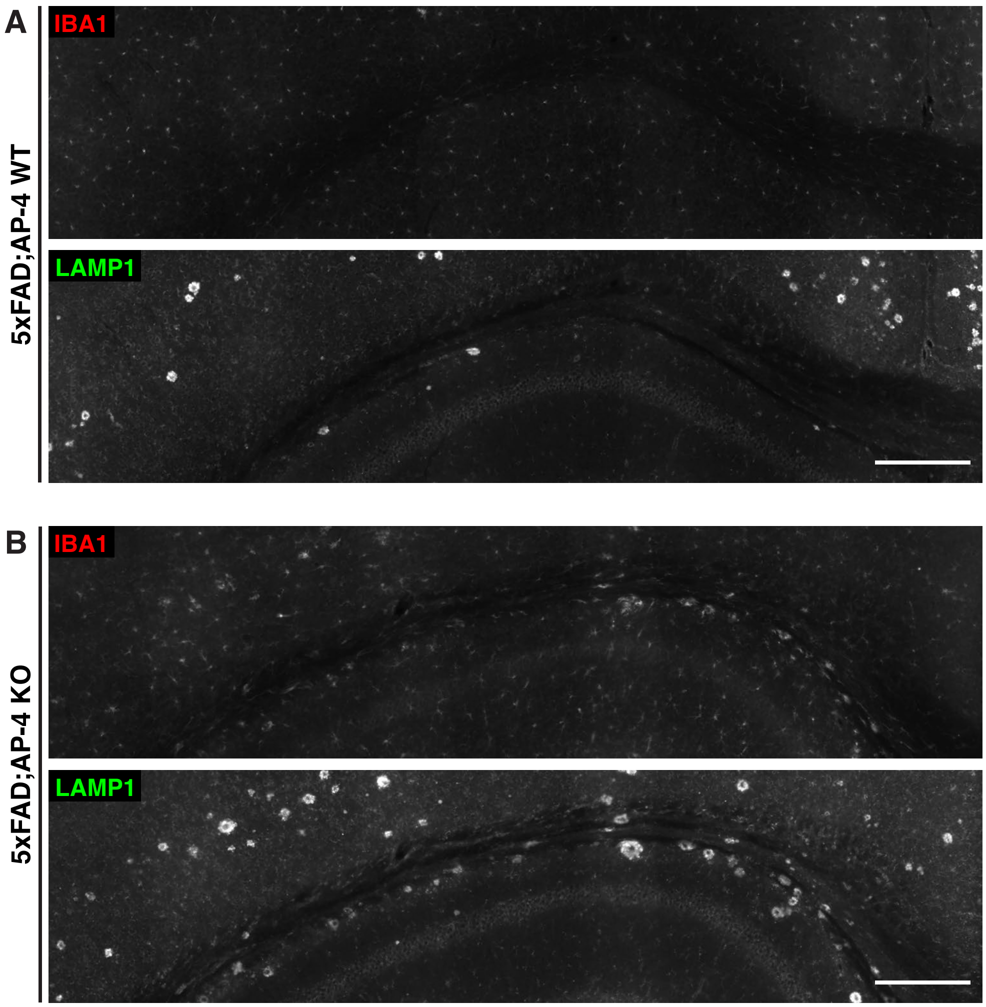


**Figure S4**

**Figure S4: Loss of AP-4 complex leads to increased gliosis in 3-month-old 5xFAD mice**

(A, B) Greyscale counterparts of the stitched images of corpus callosum and hippocampus of 3-month-old 5xFAD mice with AP-4 (WT; A) or lacking AP-4 (KO; B) from Figure 2A, depicting neuritic plaques stained with LAMP1 (green; lysosomes) and microglia stained with IBA1 (red). Bar, 100μm. Greyscale images highlight increased gliosis in the 5xFAD; AP-4 KO animals.

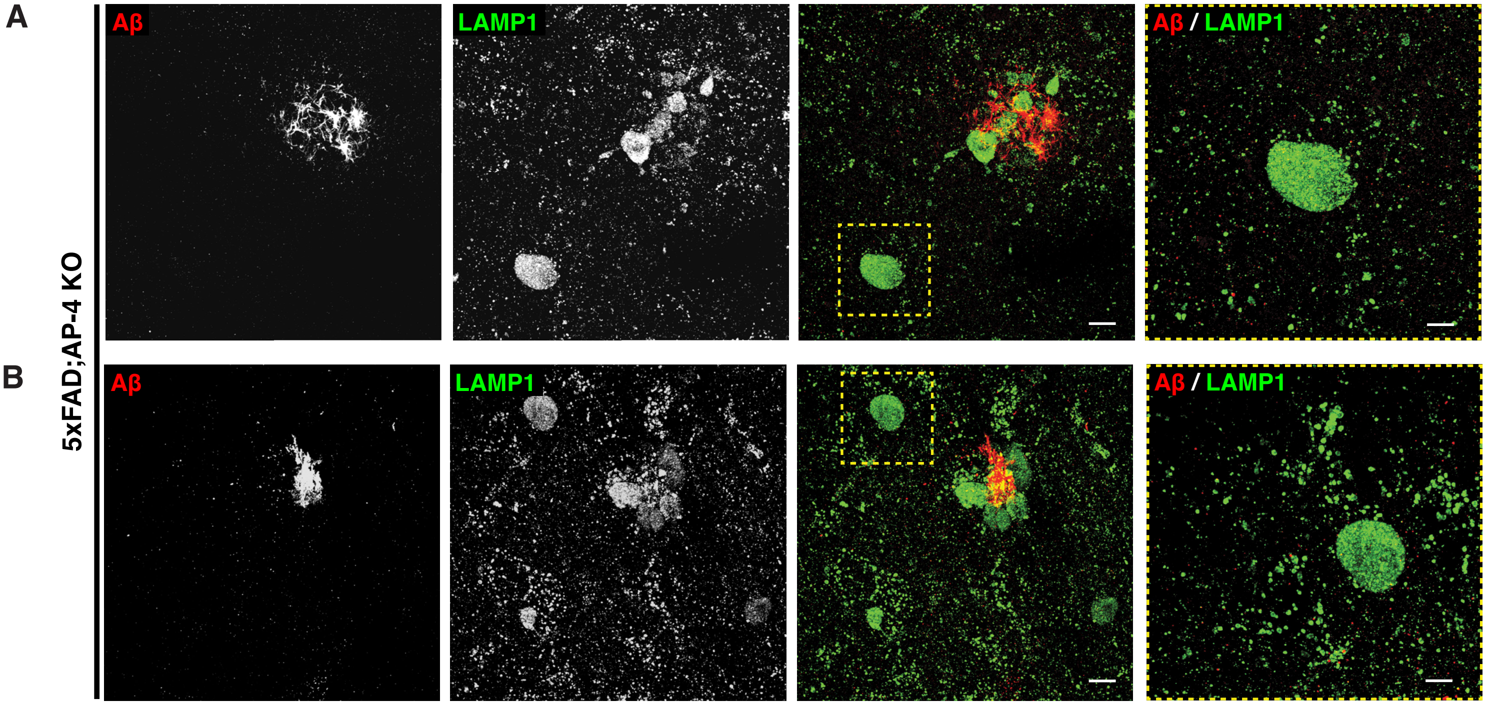


**Figure S5**

**Figure S5: Expansion microscopy images of neuritic plaques and AP-4 dystrophies in 5xFAD; AP-4 KO mice**

(A, B) Maximum intensity projection (MIP) images showing neuritic plaques and AP-4 dystrophies in the hippocampus of a 3-month-old female 5xFAD mouse lacking AP-4 (KO). The brain slices were stained for LAMP1 (green; lysosomes) and Aβ (red; amyloid aggregates) and expanded about four-fold. The dashed yellow boxes in the third column highlight the AP-4 dystrophies which are also enlarged and depicted in the fourth column. Bar, 5μm (20μm); 2.5μm (10μm) for the fourth column. Scale bars are provided at the pre-expansion scale (with the corresponding post-expansion size indicated in brackets).

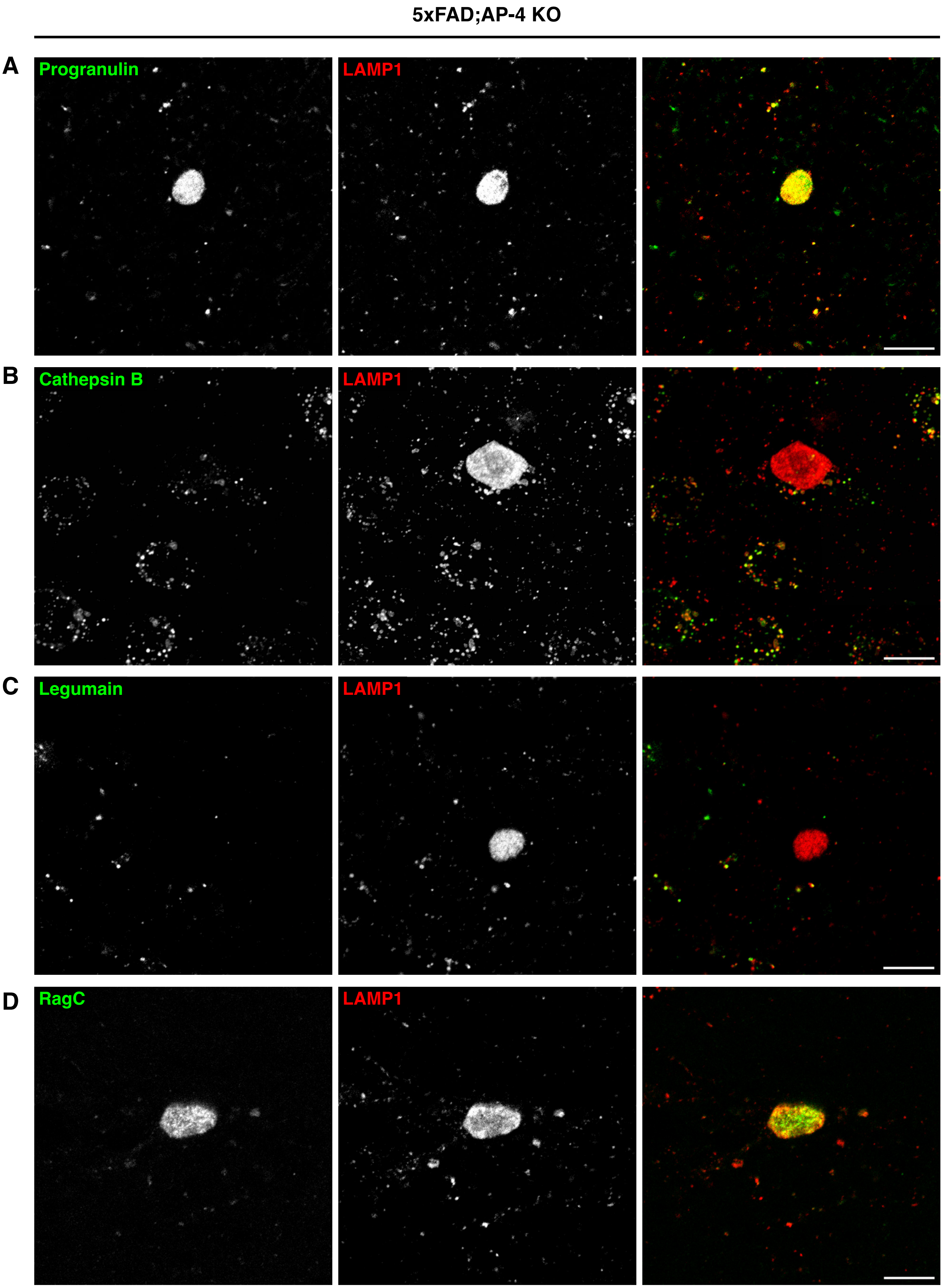
5xFAD;AP-4 KO
A
Progranulin
LAMP1
B
Cathepsin B
LAMP1
C
Legumain
LAMP1
D
RagC
LAMP1

Figure S6

**Figure S6: Expression of Cathepsin B, Legumain, Progranulin, and Rag C in AP-4 Dystrophies**

(A) High-resolution confocal images of Progranulin (green), and LAMP1 (red) enrichment in an AP-4 dystrophy in mice lacking AP-4 (AP-4ε KO mice). Bar, 10μm. (B) High-resolution confocal images of the soluble protease, Cathepsin B (green), and LAMP1 (red) in AP-4 dystrophy in 3-month-old mice lacking AP-4 (AP-4ε KO mice). Bar, 10μm. (C) High-resolution confocal images of lysosomal protease, Legumain/AEP (Asparaginyl endopeptidase) (green), and LAMP1 (red) in AP-4 dystrophy in mice lacking AP-4 (AP-4ε KO mice). Bar, 10μm. (D) High-resolution confocal images of small GTPase RagC (green), and LAMP1 (red) in AP-4 dystrophy in mice lacking AP-4 (AP-4ε KO mice). Bar, 10μm.

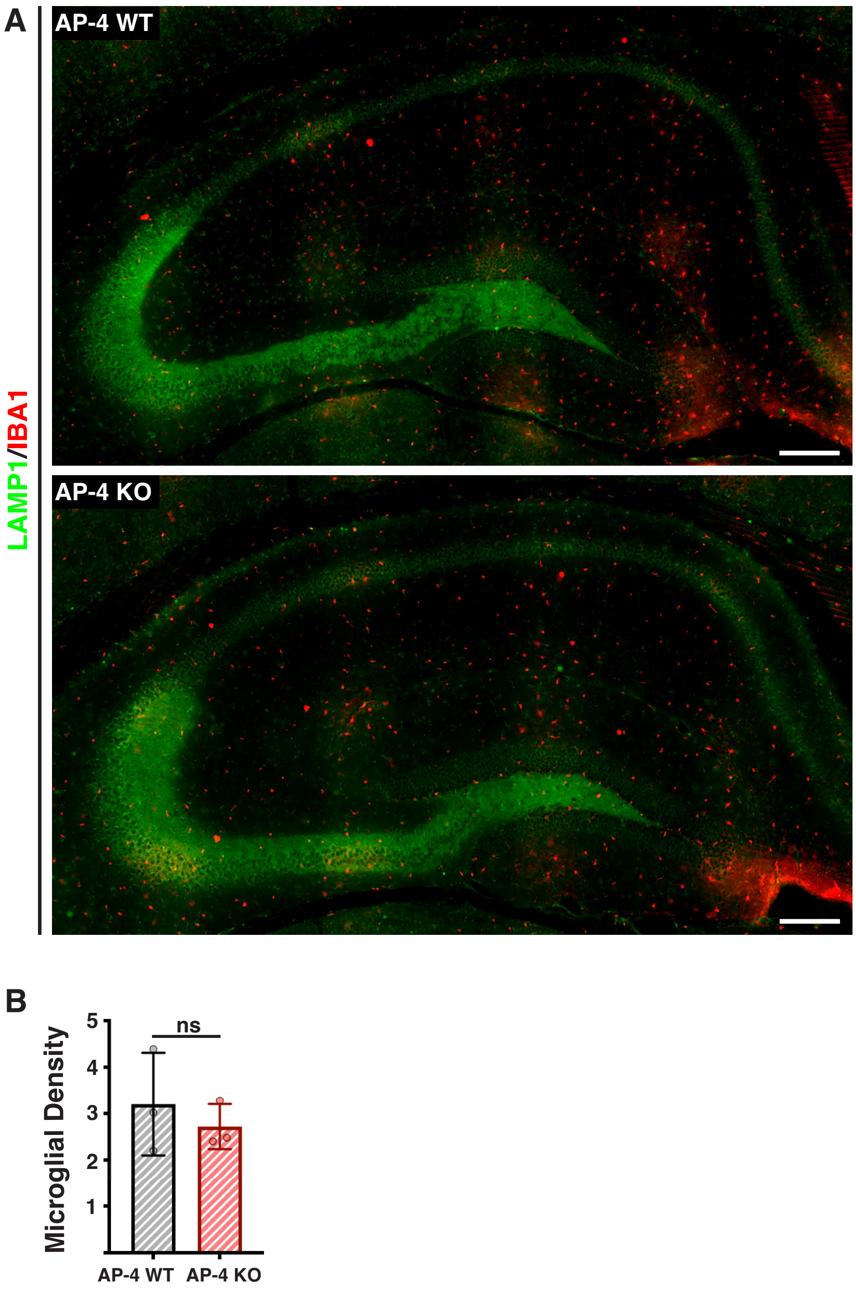
A
AP-4 WT
AP-4 KO
LAMP1/IBA1
B
Microglial Density
ns
5
4
3
2
1
AP-4 WT
AP-4 KO

Figure S7

**Figure S7: Loss of AP-4 complex alone (in absence of FAD mutations), does not increase gliosis**

(A) Stitched images of LAMP1 (green; lysosomes) and IBA1 (red; microglia) in 6–8-month-old AP-4 WT and AP-4ε KO animals. Bar, 100μm. (B) Quantification of microglial burden (number of microglia per 10,000$\mu m^2$) in the corpus callosum and hippocampus of 6-8- month-old AP-4 WT and AP-4ε KO animals (sex-matched littermates). Mean ± SEM, N = 3, ns, P = 0.5267, unpaired *t* test.

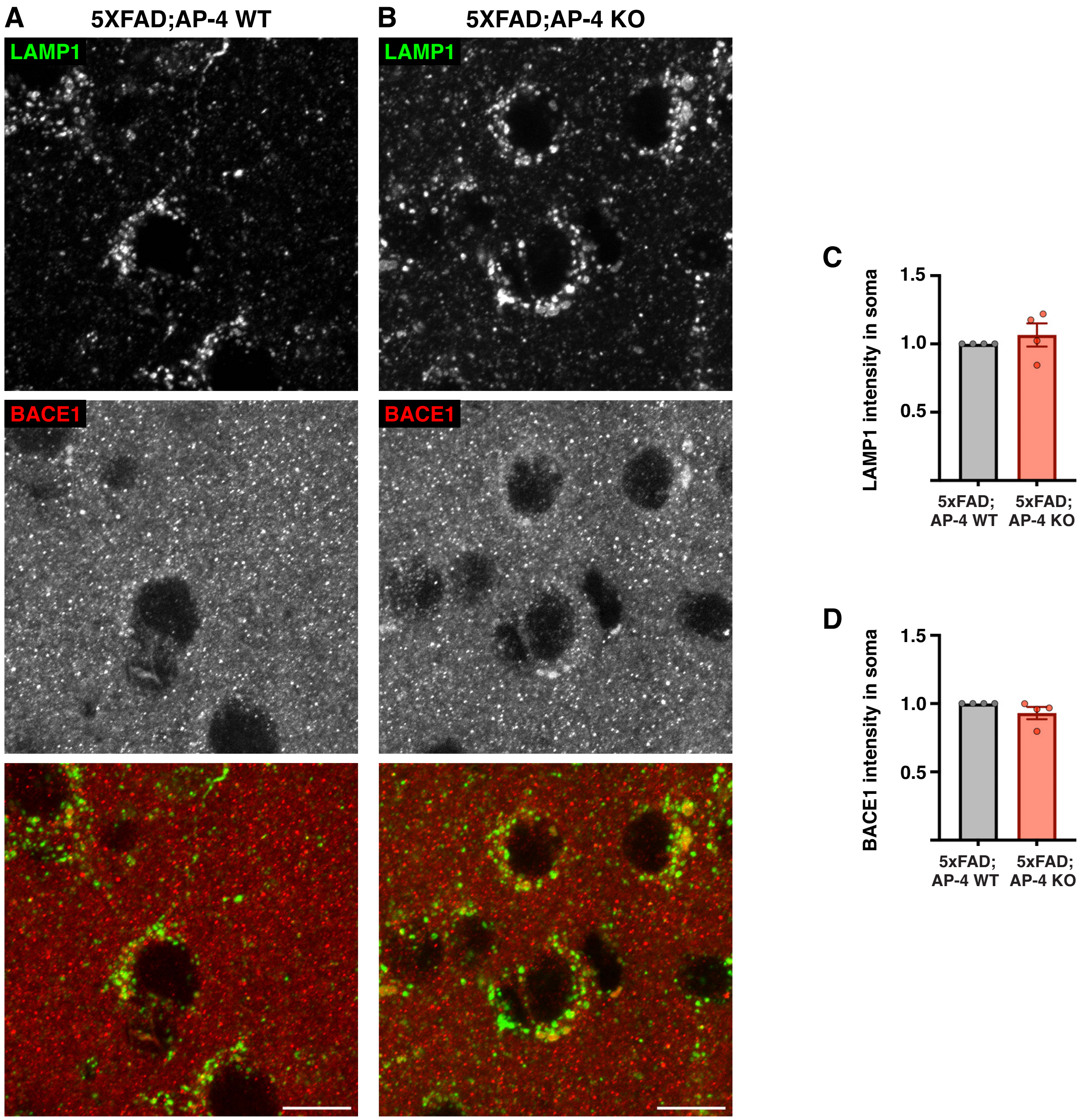
A
5XFAD;AP-4 WT
B
5XFAD;AP-4 KO
LAMP1
BACE1
LAMP1
BACE1
C
LAMP1 intensity in soma
1.5
1.0
0.5
5xFAD; AP-4 WT
5xFAD; AP-4 KO
D
BACE1 intensity in soma
1.5
1.0
0.5
5xFAD; AP-4 WT
5xFAD; AP-4 KO

**Figure S8**

**Figure S8: Analysis of LAMP1 and BACE1 intensities in lysosomes of neuronal cell bodies in 5xFAD; AP-4 WT and 5xFAD; AP-4 KO mice**

(A, B) High-resolution confocal images of LAMP1 (green; lysosomes) and BACE1 (red; BACE1-positive vesicles) in neuronal cell bodies of 3-month-old 5xFAD mice with AP-4 (WT; A) or lacking AP-4 (KO; B), (C) Quantification of fold change in mean intensity of LAMP1 in neuronal cell bodies of 3-month-old female 5xFAD mice with AP-4 (WT; Grey) or lacking AP-4 (KO; Red). Mean ± SEM, N= 4. (D) Quantification of fold change in mean intensity of BACE1 in neuronal cell bodies of 3-month-old female 5xFAD mice with AP-4 (WT; Grey) or lacking AP-4 (KO; Red). Mean ± SEM, N= 4.

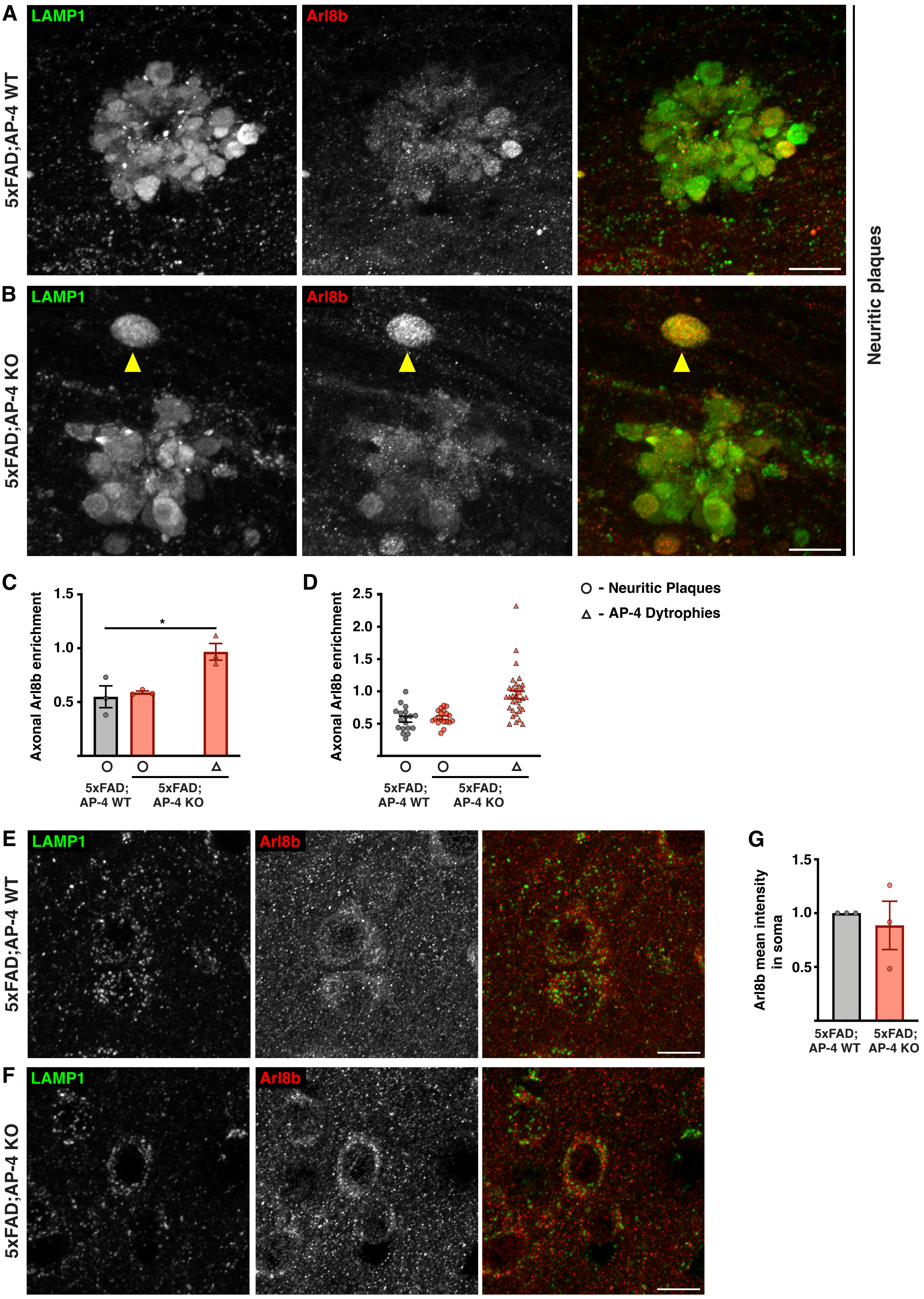
A
B
C
D
E
F
G
LAMP1
Arl8b
5xFAD;AP-4 WT
5xFAD;AP-4 KO
Neuritic plaques
Axonal Arl8b enrichment
1.5
1.0
0.5
2.5
2.0
*
5xFAD;
AP-4 WT
AP-4 KO
○ - Neuritic Plaques
△ - AP-4 Dytrophies
Arl8b mean intensity
in soma

Figure S9

**Figure S9: Arl8b is enriched in AP-4 Dystrophies**

(A, B) High-resolution confocal images of LAMP1 (green) and Arl8b (red) depicting neuritic plaques (A, B) and AP-4 dystrophies (B; yellow arrowhead) in the corpus callosum of 3-month-old 5xFAD mice with AP-4 (WT; A) or lacking AP-4 (KO; B). Note that AP-4 dystrophies are only observed in mice lacking AP-4. Bar, 10μm. (C) Quantification of mean relative enrichment of Arl8b in axonal dystrophies of neuritic plaques and AP-4 dystrophies compared to neuronal soma in 3-month-old female 5xFAD mice with AP-4 (WT; Grey) or lacking AP-4 (KO; Red). Mean ± SEM, N= 3, *, $P < .05$, one way-ANOVA with Dunnett's post-test. Neuritic plaques- circle and AP-4 dystrophies- triangle. (D) Plot depicting relative Arl8b enrichment from individual neuritic plaques and AP-4 dystrophies of the three animals from each genotype (Grey- 5xFAD; AP-4 WT; Red- 5xFAD; AP-4 KO). Neuritic plaques- circle and AP-4 dystrophies- triangle. (E, F) High-resolution confocal images of LAMP1 (green) and Arl8b (red) in neuronal cell bodies of 3-month-old 5xFAD mice with AP-4 (WT; E) or lacking AP-4 (KO; F). Bar, 10μm. (G) Quantification of fold change in mean intensity of Arl8b in neuronal cell bodies of 3-month-old female 5xFAD mice with AP-4 (WT; Grey) or lacking AP-4 (KO; Red). Mean ± SEM, N= 3.
